# Retinoblastoma binding protein 4 maintains cycling neural stem cells and prevents DNA damage and Tp53-dependent apoptosis in *rb1* mutant neural progenitors

**DOI:** 10.1101/427344

**Authors:** Laura E. Schultz-Rogers, Maira P. Almeida, Wesley A. Wierson, Marcel Kool, Maura McGrail

**Affiliations:** Department of Genetics, Development and Cell Biology, Iowa State University, Ames, IA 50011, USA; Center for Individualized Medicine, Mayo Clinic, Rochester, MN 55905, USA; Hopp Children’s Cancer Center at the NCT (KiTZ), 69120 Heidelberg, Germany; Division of Pediatric Neuro-oncology, German Cancer Research Center (DKFZ), and German Cancer Consortium (DKTK), 69120 Heidelberg, Germany

## Abstract

Retinoblastoma-binding protein 4 (Rbbp4) is a WDR adaptor protein for multiple chromatin remodelers implicated in human oncogenesis. Here we show Rbbp4 is overexpressed in zebrafish *rb1*-embryonal brain tumors and is upregulated across the spectrum of human embryonal and glial brain cancers. We demonstrate *in vivo* Rbbp4 is essential for zebrafish neurogenesis and has distinct roles in neural stem and progenitor cells. *rbbp4* mutant neural stem cells show delayed cell cycle progression and become hypertrophic. In contrast, *rbbp4* mutant neural precursors accumulate extensive DNA damage and undergo programmed cell death that is dependent on Tp53 signaling. Loss of Rbbp4 and disruption of genome integrity correlates with failure of neural precursors to initiate quiescence and transition to differentiation. *rbbp4; rb1* double mutants show that survival of neural precursors after disruption of Rb1 is dependent on Rbbp4. Elevated Rbbp4 in *Rb1-*deficient brain tumors might drive proliferation and circumvent DNA damage and Tp53-dependent apoptosis, lending support to current interest in Rbbp4 as a potential druggable target.

**Author Summary:** Examining the developmental mechanisms controlling neural stem and progenitor cell behavior is critical to our understanding of the processes driving brain tumor oncogenesis. Chromatin remodelers and their associated adaptor proteins are thought to be key drivers of brain development and disease through epigenetic regulation of gene expression and maintenance of genome integrity, but knowledge of their *in vivo* roles in vertebrate neurogenesis is limited. The chromatin remodeler adaptor protein Rbbp4 has recently been shown to function in a mouse model of neuroblastoma and in glioblastoma multiforme cell resistance to the chemotherapeutic temozolomide. However, an *in vivo* requirement for Rbbp4 in neurogenesis has only just been shown by isolation of a recessive lethal mutation in zebrafish *rbbp4*. Here we provide conclusive genetic evidence that zebrafish rbbp4 is essential in neural stem and progenitor cell function during development. Our data reveal for the first time *in vivo* that Rbbp4 prevents DNA damage and activation of Tp53 signaling pathway that leads to programmed cell death. Importantly, neural progenitors that are mutant for the tumor suppressor Rb1 also depend on Rbbp4 for survival. Finally, we show that neural stem cells that have lost Rbbp4 cease dividing, and may enter a senescent like state. Together, these observations provide novel evidence that elevated expression of Rbbp4 in *rb1-*mutant tumors may contribute to cancer cell survival by blocking senescence and/or DNA damage-induced cell death.

## Introduction

The Retinoblastoma Binding Protein 4 (RBBP4) is a WD40 Repeat (WDR) protein that functions as a chromatin assembly factor and adaptor for multiple chromatin remodelers regulating gene expression and DNA repair. RBBP4 associates with chromatin through histone H3 and H4 binding sites on the surface and side of the β-propeller, respectively (1–3). These locations are potential sites for chemical inhibitors of RBBP4 activity (4). RBBP4 is a component of the H3K27methyltransferase polycomb repressive complex PRC2 (2) and the nucleosome remodeling and histone deacetylase complex NuRD (5–7) and binds the histone acetyltransferase p300/CREB binding protein complex (8). RBBP4-dependent recruitment of chromatin regulators has recently been shown to control glioblastoma multiforme cell resistance to temozolomide through p300 activation of DNA repair pathway genes (9), and tumor progression in neuroblastoma xenografts by PRC2 silencing of tumor suppressors (10). Only recently has an *in vivo* requirement for RBBP4 in vertebrate development and neurogenesis been demonstrated by mutational analysis in zebrafish (11). Given the importance of understanding how developmental mechanisms contribute to the cellular and molecular basis of cancer (12), a more detailed examination of the role of RBBP4 in neural development would provide additional knowledge into its possible oncogenic roles in brain tumors.

RBBP4 was first discovered as a binding partner of the tumor suppressor RB1 in yeast (13) and studies in *C.elegans* revealed the RBBP4 homolog *lin-53* negatively regulates vulval precursor cell fate by repression of genes required for vulval induction (14). Drosophila and human RBBP4 were shown to be a component of the RB1-like DREAM complex, which binds E2F target gene promoters to repress cell cycle gene expression and promote entry into quiescence (15). These early *in vivo* studies indicated RBBP4 functions in cell cycle progression and cell fate specification through transcriptional regulation by RB family members. RB1 has been well studied in mouse and shown to be essential for embryonic and adult neurogenesis (16, 17). Zebrafish *rb1* is also an essential gene required for neural development (18, 19). Mutant analysis plus live brain imaging revealed zebrafish *rb1* mutant neural progenitors reenter the cell cycle and fail to progress through mitosis, whereas loss of *rbbp4* induces neural progenitor apoptosis (11). Together these studies suggest Rbbp4 and Rb1 cooperate to ensure neural progenitor cell cycle exit and differentiation.

In this study we use developmental genetics to further investigate the *in vivo* role of Rbbp4 in zebrafish neurogenesis and examine the interaction of Rbbp4 and Rb1 in neural progenitor cell fate. Rbbp4 is enriched in zebrafish neural stem and precursor cells during development and in adult brain. Rbbp4 is also upregulated across many human embryonal and glial brain tumors. We find that *rbbp4* mutant neural stem cells become hypertrophic, while neural precursors accumulate DNA damage and undergo Tp53-dependent apoptosis. *in situ* analysis demonstrates *rbbp4* mutant progenitors are lost before exiting the cell cycle and initiating the G0 gene expression program. Double mutant analysis shows Rbbp4 is required for the survival of *rb1*-mutant neural progenitors. Our developmental analyses suggest a potential oncogenic role for Rbbp4 in *rb1*-defective brain cancer cells by promoting proliferation and preventing DNA damage induced apoptosis.

## Results

### Rbbp4 protein is expressed in zebrafish adult neural stem/progenitors and in *rb1*-embryonal brain tumors

We previously showed by RNA-Seq and qRT-PCR that *rbbp4* is overexpressed in zebrafish *rb1-*brain tumors, and demonstrated it is required in neural development by isolating a larval lethal homozygous mutant allele *rbbp^Δ4is60^* (11). Here we used a commercially available antibody to examine Rbbp4 localization in zebrafish tissues. Western blotting with the anti-RBBP4 polyclonal antibody detects a single band of ~47 kDa in whole protein extract from 5 day post fertilization (dpf) wildtype zebrafish larvae that is absent in *rbbp4^Δ4/Δ4^* homozygous mutant extract (Fig S1A). In cross sections of 5 dpf head tissue Rbbp4 labeled nuclei are detected throughout the brain, but the labeling was again absent in *rbbp4^Δ4/Δ4^* homozygous mutant tissue (Fig S1B, S1C), confirming the specificity of the antibody. In wild type adult zebrafish, stem and progenitor cells line the ventricle and subventricular zone of midbrain structures (Fig 1A). Rbbp4 is detected in cells lining the midbrain ventricle where stem cells reside (Fig 1B-D) and is enriched in the nuclei of cells with high levels of proliferating cell marker PCNA (Fig 1D arrow). Rbbp4 was also present in nuclei of cells throughout the midbrain peri-granular zone (Fig 1E). In *rb1-*brain tumor tissue (Fig 1F) high levels of nuclear Rbbp4 were detected in cells throughout the lesion (Fig 1G) that frequently co-labeled with PCNA (Fig 1H, 1I). These results confirm overexpression of Rbbp4 protein in zebrafish *rb1-*brain tumors, as predicted by our previous RNA-Seq analysis (11).

**Fig 1.**
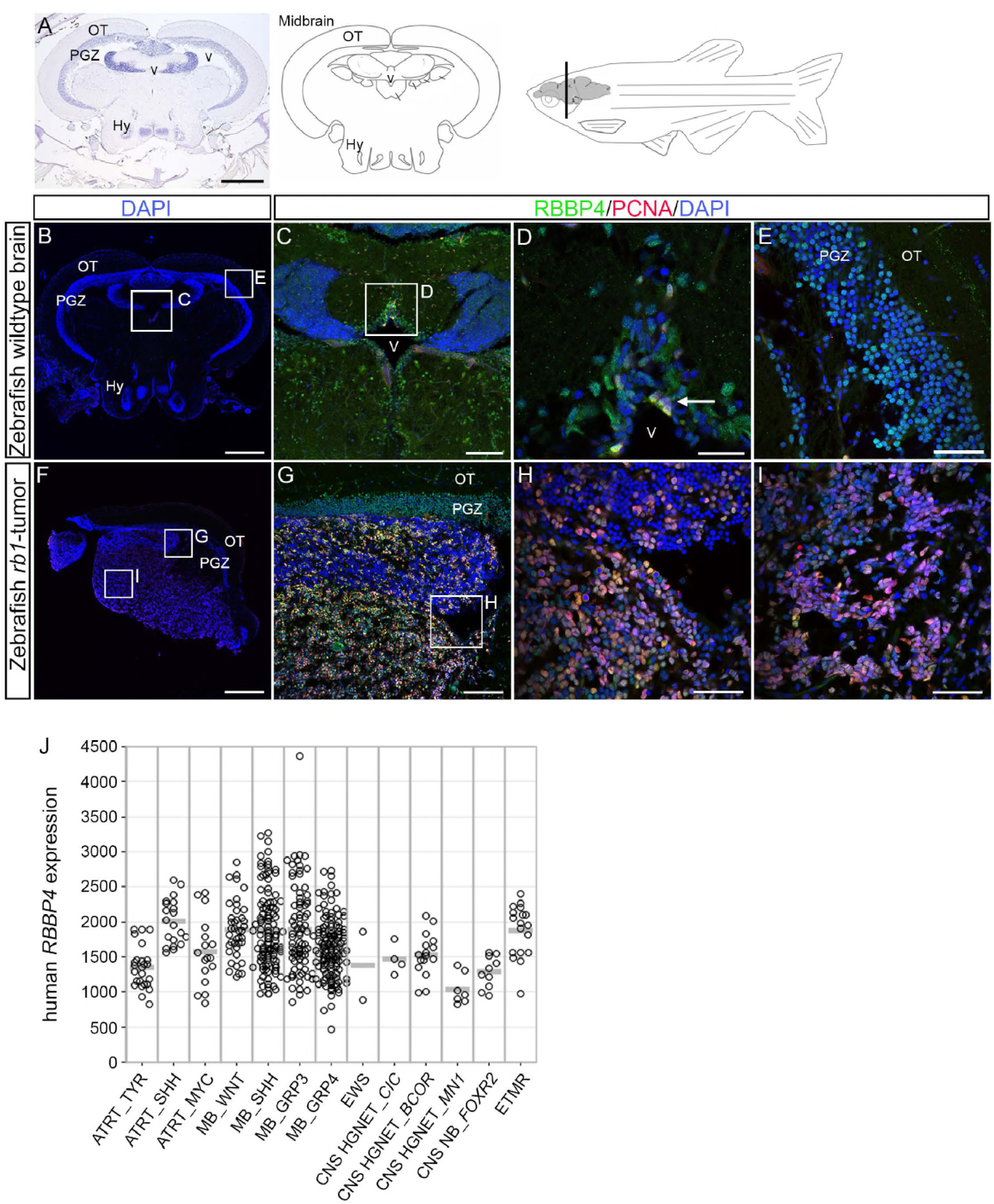
Rbbp4 is localized in the nucleus in adult neural stem and progenitors and is upregulated in zebrafish *rb1* and human embryonal brain tumors. (A) Hematoxylin-labeled cross-section and diagram of adult zebrafish midbrain. Representative transverse cryosections (n=3 independent animals) through adult zebrafish midbrain (B-E) or *rb1-*embryonal brain tumor (F-I) labeled with antibodies to Rbbp4 (green), proliferation maker PCNA (red), and nuclear stain DAPI (blue). Rbbp4 is detected in the nuclei of cells lining the midbrain ventricle (C, D) and throughout the peri-granular zone (E), and is enriched in PCNA positive cells (D, arrow). Section through *rb1-*brain tumor shows expansion of the ventricular zone (F, G). The lesion is filled with Rbbp4 positive cells that frequently co-label with PCNA (H, I). Hy, hypothalamus; OT, optic tectum; PGZ, peri-granular zone; v, midbrain ventricle. Scale bars: A, E 400 μm; B, F 100 μm; D, G, H 40 μm; C 20 μm. See also Fig S1. (J) *RBBP4* expression in human embryonal tumors. ATRT, Atypical Teratoid/Rhabdoid Tumors: ATRT_TYR (n=25), ATRT_SHH (n=20), ATRT_MYC (n=17); MB, Medulloblastoma: MB_WNT (n=40), MB_SHH (n=112), MB_GRP3 (n=81), MB_GRP4 (n=135); EWS, Ewing Sarcoma (n=2); CNS EFT_*CIC*, CNS Ewing sarcoma family tumor with *CIC* alteration (n=4); CNS HGNET_*BCOR*, CNS high-grade neuroepithelial tumor with *BCOR* alteration (n=16); CNS HGNET_*MN1*, CNS high-grade neuroepithelial tumor with *MN1* alteration (n=7); CNS-NB-*FOXR2*, CNS neuroblastoma with *FOXR2* activation (n=10); EMTR, embryonal tumors with multilayered rosettes (n=18).

### Human RBBP4 is upregulated across the spectrum of aggressive brain cancers

We examined expression data from 2284 human brain tumor samples (German Cancer Research Center) to determine if *RBBP4* is upregulated in human embryonal tumors and other aggressive brain cancers. Molecular profiling has classified human embryonal central nervous system primitive neuroectodermal tumors (CNS-PNETs) into major groups with distinct molecular subtypes (20–22). *RBBP4* was upregulated across all embryonal tumors and subtypes (Fig 1J) as well as in ependymal, glial, oligodendroglial and astrocytic tumors (Fig S1D). Together these results indicate elevated *RBBP4* may be a common signature in aggressive central nervous system tumors.

### Zebrafish Rbbp4 protein is ubiquitously localized in embryonic neuroblasts and varies in neural cell populations in the larval midbrain

Rbbp4 expression was examined in the 2 dpf embryonic (Fig 2A, 2B) and 5 dpf larval (Fig 2C, 2D) wildtype zebrafish developing brain. At 2 dpf nuclear Rbbp4 is present in all neural precursor cells throughout the midbrain and retina and co-labels with proliferating cell marker PCNA (Fig 2E, 2F, S2). Rbbp4 and PCNA were highly enriched in neural stem cells lining the midbrain ventricle (Fig 2G) and at the ciliary marginal zone (CMZ) of the retina (Fig 2H). The end of 2 dpf marks the transition from embryonic to larval neurogenesis and growth in the brain becomes asymmetric (23, 24). By 5 dpf, proliferative zones are confined to the midbrain ventricles and dorsal tectum and the retinal ciliary marginal zone (25, 26). In the 5 dpf midbrain (Fig 5I) Rbbp4 is significantly reduced or absent from most cells in the proliferative zones (Fig 2J, 2K and 2H outlined areas; Fig S3) but is detectable in PCNA-positive cells (Fig 2K and 2L arrows; Fig S3). Varying levels of Rbbp4 were present in cells adjacent to the ventricle and in neurons throughout the parenchyma. At 5 pdf the retina is laminated into 3 distinct nuclear layers and stem cells and progenitors are restricted to the ciliary marginal zone (Fig 2M). Rbbp4 is detectable in PCNA-positive cells at the retinal ciliary marginal zone (Fig 2N and 2O outlined area, Fig S3). In the central retina, high levels of Rbbp4 were restricted to neurons in the ganglion cell layer and at the vitreal side of the inner nuclear layer (Fig 2P, Fig S3). A low level of Rbbp4 was detected in outer nuclear layer photoreceptors (Fig 2P). In summary, in the larval midbrain and retina Rbbp4 is present in proliferating stem and progenitor cells and is elevated to varying levels in other neuronal cell populations. This suggests a role for Rbbp4 in both neural stem and precursor cells during brain development.

**Fig 2.**
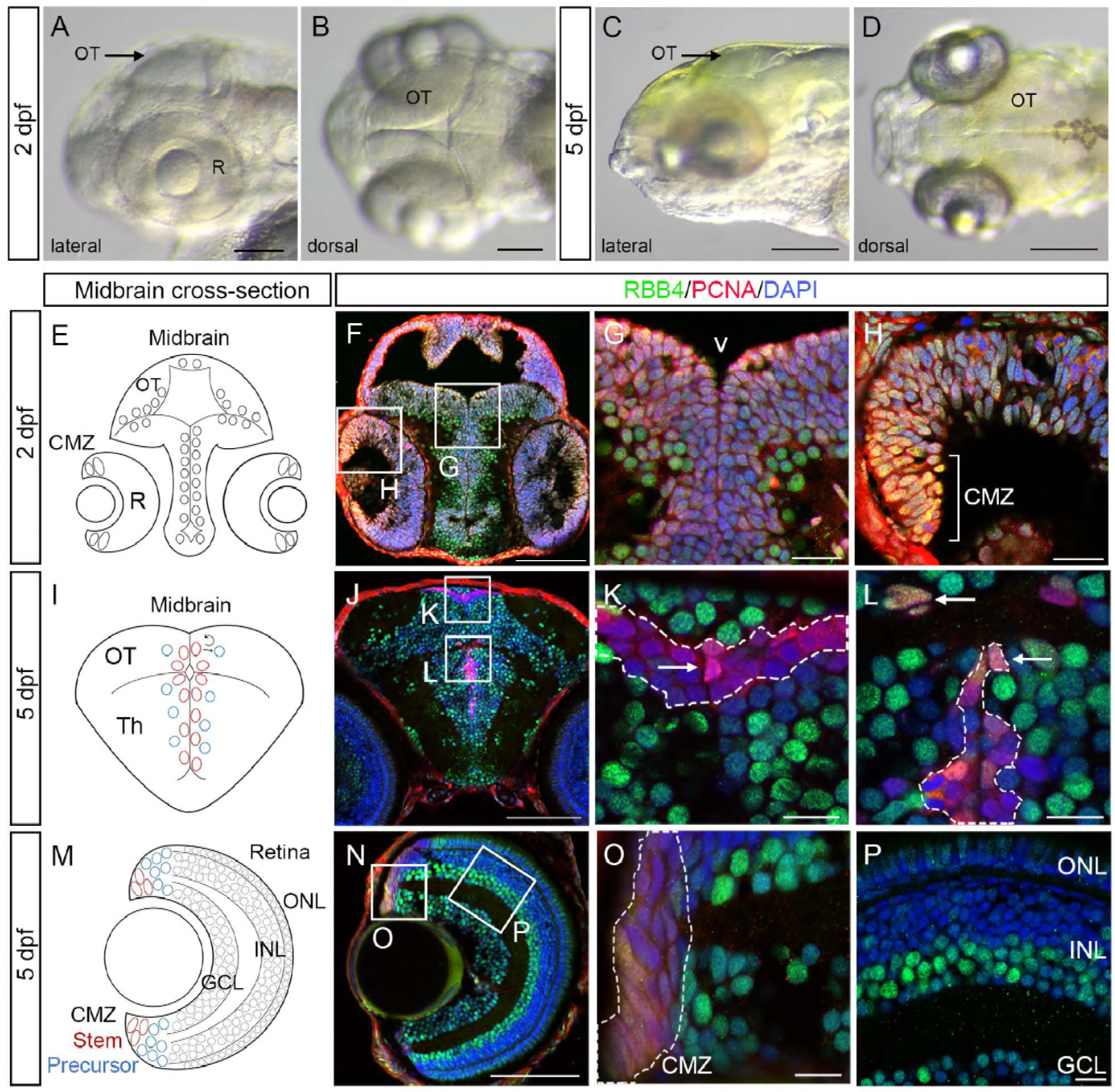
Rbbp4 is highly expressed in embryonic neuroblasts and shows variable levels in neural stem cells, precursors and neurons in the larval brain. Lateral and dorsal views of head region in zebrafish 2 dpf embryo (A, B) and 5 dpf larva (C-D) with midbrain optic tectum (OT, arrows) and retina (R) labeled. (E-P) Representative transverse cryosections (n=3) through 2 dpf and 5dpf wildtype zebrafish midbrain labeled with antibodies to Rbbp4 (green), proliferation maker PCNA (red), and nuclear stain DAPI (blue). (E) Diagram of location of stem cells lining the midbrain ventricle and at the ciliary marginal zone (CMZ) of the retina. (F-H) Boxes outline the ventricle at the top of the thalamus in the midbrain (G) and the retina ciliary marginal zone (H, bracket). See also Fig S1. (I) Diagram of 5 dpf midbrain showing location of stem cells (red) and progenitors (blue) at the ventricle in the dorsal optic tectum and ventral thalamus (Th). (J-L) 5 dpf wildtype zebrafish midbrain. Higher magnification view of the medial tectal proliferative zone (K) and midbrain ventricle (L). (M) Diagram of 5 dpf retina showing location of stem cells and retinal precursors at the ciliary marginal zone and the three nuclear retinal layers. (N-P) 5 dpf retina. Higher magnification view of the ciliary marginal zone (O) and central laminated layers of the retina (P). The highly proliferative, PCNA+ population of cells are outlined; arrows indicate cells co-labeled with Rbbp4 and PCNA. Rbbp4 is enriched in a subset of neurons in the midbrain and vitreal side of the retinal inner nuclear layer. See also Fig S2. GCL, ganglion cell layer; INL, inner nuclear layer; ONL, outer nuclear layer. Scale bars: 100 μm A, B, F, J, N; 200 μm C, D; 20 μm G, H; 10 μm K, L, O, P.

### Rbbp4 is essential for development of brain, eye and neural crest derived structures

Zebrafish larvae homozygous mutant for the 4 base pair frameshift mutation *rbbp4^Δ4^* display severe defects in neurogenesis, presenting with microcephaly and microphthalmia (Fig 3A-3D). Alcian blue staining revealed several abnormalities in formation of head cartilage structures (Fig 3E-3H), including the ceratohyal cartilage (ch), ceratobranchial cartilage (cbs), and Meckel’s cartilage (m). These results suggest that Rbbp4 is necessary for development of the central nervous system and neural crest derived structures. Ubiquitous overexpression of *rbbp4* cDNA by a *Tol2<ubi:rbbp4-2AGFP>* transgenic line (Fig 3I) was tested for the ability to rescue the *rbbp4^Δ4/Δ4^* phenotype. The transgene did not affect development in wild type embryos or viability and fertility in adults (Fig 3J, Table S1) and was able to rescue the gross morphological defects in *rbbp4^Δ4/Δ4^* mutants (Fig 3K and Fig S3). Midbrain height and eye width measurements in *Tol2<ubi:rbbp4-2AGFP>* and *rbbp4^Δ4/Δ4^;Tol2<ubi:rbbp4-2AGFP>* transgenic embryos confirmed rescued animals showed no significant difference in morphology from wildtype (Fig 3L and 3M). *rbbp4^Δ4/Δ4^; Tg*(*Tol2<ubi:rbbp4-2AGFP>*) individuals are viable and become fertile adults. These results demonstrate Rbbp4 is essential for larval viability, midbrain and retina development, and formation of neural crest derived cartilage.

**Fig 3.**
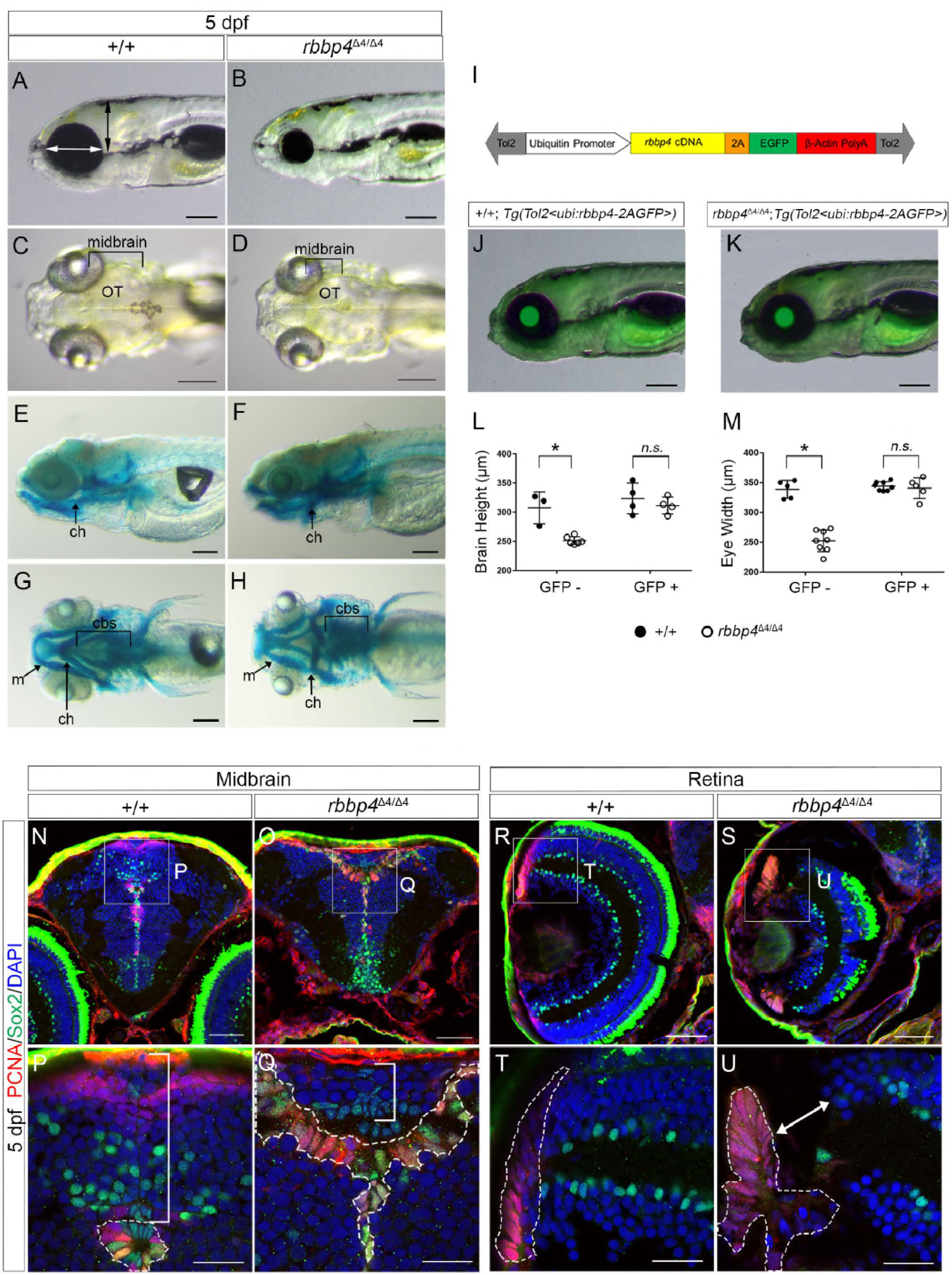
Rbbp4 is required for zebrafish brain and neural crest development and persistent neurogenesis in the midbrain optic tectum and retina. (A-K) *rbbp4* cDNA rescue of *rbbp4^Δ4/Δ4^* homozygotes demonstrates requirement for Rbbp4 in brain and neural crest development. (A) Lateral view of 5 dpf wildtype zebrafish indicating location of measurements for head height (black arrow) and eye width (white arrow). (B) Lateral view of 5 dpf *rbbp4^Δ4/Δ4^* homozygote showing gross defects including microcephaly and microphthalmia. Lateral and ventral views of 5 dpf wildtype (C, E) and *rbbp4^Δ4/Δ4^* homozygous (D, F) larva stained with alcian blue to reveal cartilage structures. (G) Schematic of *Tol2<ubi:rbbp4–2AGFP>* cDNA rescue construct driving expression of *rbbp4–2AGFP* by the *ubiquitin* promoter. (H) 5 dpf transgenic *Tg*(*Tol2<ubi:rbbp4-2AGFP>*) and (I) 5 dpf transgenic *rbbp4^Δ4/Δ4^*; *Tg*(*Tol2<ubi:rbbp4-2AGFP>*) larva showing rescue of the *rbbp4^Δ4/Δ4^* homozygous mutant phenotype. (J) *rbbp4^Δ4/Δ4^* homozygotes (n=6) have a significantly smaller head than wildtype (n=3, *p*=0.0015). Transgenic GFP+ *rbbp4^Δ4/Δ4^* homozygotes (n=4) show no significant difference in head height compared to GFP+ +/+ wildtype (n=4) (p=0.4595). (K) *rbbp4^Δ4/Δ4^* homozygotes (n=8) have a significantly smaller eye than wildtype (n=5, *p*=0.0001). Transgenic *rbbp4^Δ4/Δ4^* GFP+ (n=5) show no significant difference in eye size compared to GFP+ +/+ wildtype (n=8, *p*=0.6293). Data represent mean ± s.d. *p* values calculated with one-tailed Student’s t-test. ch, ceratohyal cartilage; m, Meckel’s cartilage; cbs, ceratobranchial cartilage. See also Fig S2 and Table S1. (L-S) Transverse sections of midbrain and retina from wild type +/+ (n=5) and homozygous mutant *rbbp4^Δ4/Δ4^* (n=4) 5 dpf zebrafish larvae labeled with antibodies to proliferative marker PCNA (red) and stem/progenitor marker Sox2 (green) and nuclear stain DAPI (blue). (L, N) In wildtype PCNA and Sox2-positive stem cells line the midbrain ventricle (L, N dashed outline). Sox2-positive progenitors lie adjacent to the ventricle in the tectum (N bracket). (N, O) PCNA and Sox2-positive cells with enlarged morphology line the midbrain ventricles in *rbbp4^Δ4/Δ4^* mutant (O outline). The dorsal tectum is significantly reduced in height in the *rbbp4^Δ4/Δ4^* mutant (O bracket). (P, R) In wild type retina PCNA-positive stem cells are restricted to the outer edge of the ciliary marginal zone (R outline). Sox2-positive amacrine and displaced amacrine cells are distributed throughout the inner nuclear and ganglion cell layers. (Q, S) PCNA-positive stem cells with enlarged, hypertrophic morphology at the ciliary marginal zone in *rbbp4^Δ4/Δ4^* mutant retina (S outline). Arrow shows missing tissue adjacent to the ciliary marginal zone. Sox2-positive cells representing early born amacrine cells are present in the retinal layers. See also Fig S3. Scale bars: A-F 100 μm; 50 μm L, M, P, Q; 20 μm N, O, R, S.

### Loss of Rbbp4 leads to stem cell hypertrophy and defects in midbrain neurogenesis

The effect of loss of Rbbp4 on specific neural cell populations in the 5 dpf larval midbrain and retina was examined with proliferation, stem, neural, and glial markers (Fig 3N-3U, Fig S5). In wildtype (Fig 3N) and *rbbp4^Δ4/Δ4^* mutant (Fig 3O) the stem/progenitor marker Sox2 was present in cells lining the midbrain ventricle and adjacent to the ventral thalamus. Sox2 positive cells were also scattered throughout the wild type dorsal tectum (Fig 3P bracket). The dorsal tectum was dramatically reduced in *rbbp4^Δ4/Δ4^* and only low levels of Sox2 were detected (Fig 3Q bracket). Sox2-positive cells frequently co-labeled with PCNA at the midbrain ventricle in wildtype (Fig 3P outline) and *rbbp4^Δ4/Δ4^* (Fig 3Q outline), however, the *rbbp4^Δ4/Δ4^* cells showed an enlarged, hypertrophic morphology. In the retina, high levels of PCNA labeling was present in stem cells at the ciliary marginal zone in both wildtype (Fig 3R) and *rbbp4^Δ4/Δ4^* (Fig 3S), but the *rbbp4^Δ4/Δ4^* ciliary marginal zone was expanded and stem cells were enlarged and hypertrophic (Fig 3T and 3U outlines), similar to cells in the midbrain ventricle. A gap of missing tissue separated the ciliary marginal zone and the neural retina in the *rbbp4^Δ4/Δ4^* mutant (Fig 3U arrow). These results suggest loss of Rbbp4 prevents brain and retina growth by disrupting production and survival of neural precursors. Examination of the glial markers Gfap and Blbp and neuronal markers synaptic vesicle Sv2 and Calretinin (Fig S5) revealed early neurogenesis occurred normally in the *rbbp4^Δ4/Δ4^* mutant brain and retina. Maternal Rbbp4 may protect the earliest born embryonic neural precursors, allowing them to migrate to their final location and differentiate into mature neurons and glia in the brain and retina. Together, these results indicate Rbbp4 is necessary for persistent production of neural precursors and their transition to newborn neurons and glia.

### Rbbp4 is required to prevent persistent DNA damage and apoptosis in neural precursor cells

To determine if the defect in neurogenesis in 5 dpf *rbbp4*^Δ4^*^/^*^Δ4^ mutant larva was due to activation of programmed cell death earlier in the developing brain, we examined gross morphology, apoptosis, and DNA damage in wildtype and *rbbp4*^Δ4^*^/^*^Δ4^ brain and retina at 2 and 3 dpf. Compared to wildtype, at 2 and 3 dpf the size of the optic tectum (Fig 4 A-C, arrows, OT) and midbrain-hindbrain structures (Fig E-H, brackets) was reduced in *rbbp4*^Δ4^*^/^*^Δ4^ mutant larvae. Apoptosis was examined by labeling sectioned tissue with an antibody to activated Caspase-3 and comparing the number of labeled cells between wildtype and *rbbp4*^Δ4^*^/^*^Δ4^ mutants (Fig 4 I-P). At 2 dpf in wildtype very few cells were labeled with activated Caspase-3, whereas labeling was significantly increased and present throughout the *rbbp4*^Δ4^*^/^*^Δ4^ tectum and overlapped with cells containing fragmented nuclei throughout the retinal ganglion and inner nuclear layers (Fig 4I-4L). At 3 dpf, few cells again were labeled in wildtype (Fig 4M). In *rbbp4*^Δ4^*^/^*^Δ4^, the amount of Caspase-3 labeling was reduced and was restricted to the proliferative zones in the dorsal tectum and periphery of the retina (Fig 4N). Quantification of Caspase-3 labeled cells revealed the increase in the *rbbp4*^Δ4^*^/^*^Δ4^ mutant at 2 dpf was significant in the brain and retina (Fig 4O). At 3 dpf the difference was less significant (Fig 4P), possibly due to overall reduction in cell number from programmed cell death or a decrease in production of new neural precursors. Activated Caspase-3 positive cells at the retina periphery did not co-label with neuronal marker HuCID, and nuclei appeared fragmented or pyknotic (Fig S6). This suggested that *rbbp4*^Δ4^*^/^*^Δ4^ mutant neural precursors undergo apoptosis before initiating differentiation. To examine whether apoptosis was linked to the DNA damage response, wildtype and *rbbp4*^Δ4^*^/^*^Δ4^ mutant tissues were labeled with an antibody to γ-H2AX (Fig 4Q-4X), which recognizes phosphorylated histone H2AX at DNA double strand breaks and more broadly labels damaged chromatin in nuclei in apoptotic cells (27). Similar to activated Capsase-3, there is little γ-H2AX labeling in the wildtype midbrain or retina at 2 dpf (Fig 4Q, 4S) or 3 dpf (Fig 4U), whereas there is a significant increase of γ-H2AX positive cells in *rbbp4*^Δ4/Δ4^ (Fig 4R, 4T, 4V-X). Close examination of γ-H2AX labeling in the retina at 2 dpf revealed three distinct patterns in *rbbp4*^Δ4/Δ4^ mutant cells. Some cells contained nuclear foci typical of DNA double strand breaks (Fig 4T, dotted arrow), while others had a defined nuclear ring (Fig 4L, open arrow) or pan-nuclear labeling (Fig 4T, closed arrow) indicative of extensive genome wide DNA damage. Cells with γ-H2AX rings or pan-nuclear labeling contained small fragmented nuclei characteristic of later stages of apoptosis (Fig 4T). At 3dpf γ-H2AX labeling was restricted to the proliferative zones in the *rbbp4*^Δ4/Δ4^ tectum and retina (Fig 4V). Together, the activated Caspase-3 and γ-H2AX labeling indicate failure of *rbbp4*^Δ4/Δ4^ mutant neural precursors to survive is due to persistent DNA damage and activation of programmed cell death.

**Fig 4.**
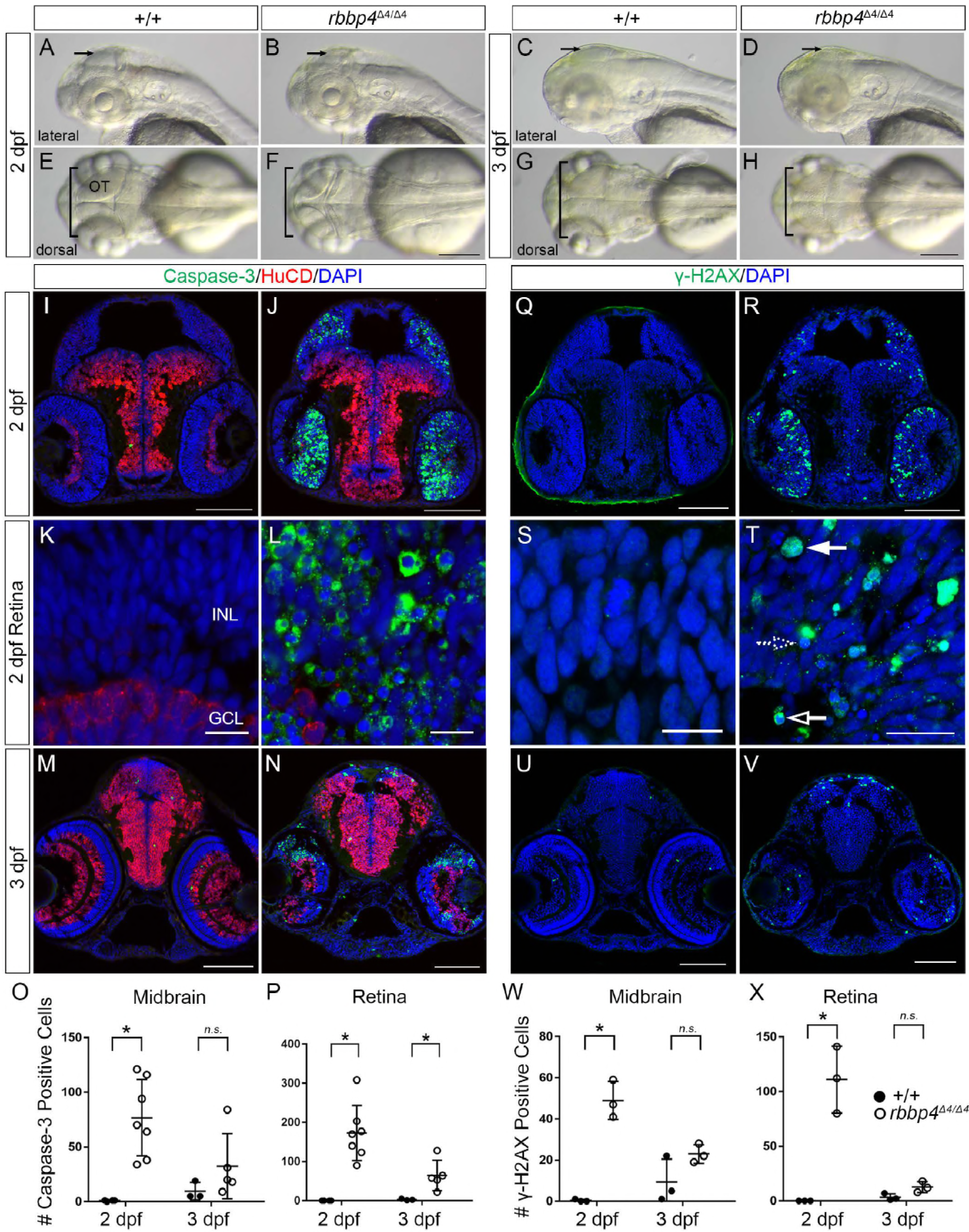
Accumulation of DNA damage and apoptosis in neural precursors underlies defects in *rbbp4^Δ4/Δ4^* midbrain and retina neurogenesis. See also Fig S3. Lateral (A-D) and dorsal (E-H) images of 2 dpf and 3 dpf larva show the size of the optic tectum (arrows, OT) and midbrain overall (brackets) is reduced in *rbbp4^Δ4/Δ4^* mutants compared to +/+ wildtype. (I-N) Transverse sections of zebrafish midbrain and retina labeled with antibodies to the apoptosis marker activated-Caspase-3 (green), neural marker HuC/D (red), and nuclear label DAPI (blue). (I, K, M) Few activated Caspase-3 labeled cells are detected in wildtype at 2dpf (n=4) and 3 dpf (n=3). (J, L, N) Extensive activated Caspase-3 labeling is present throughout the midbrain tectum and retina in the *rbbp4^Δ4/Δ4^* mutant at 2 dpf (n=7) and 3 dpf (n=5). (L) Higher magnification view of nuclear morphology shows pyknosis and nuclear fragmentation in the *rbbp4^Δ4/Δ4^* mutant 2 dpf retina. (O, P) Counts of activated Caspase-3 positive cells plotted for individual larva show a significant difference between wildtype and *rbbp4^Δ4/Δ4^* mutant at 2 dpf in the midbrain (O, *p*=0.0022) and in the retina (P, *p*=0.0010). At 3 dpf the difference is no longer significant in the midbrain (O, *p*=0.2569) and only slightly significant in the retina (P, *p*=0.0358), due to loss of cells by apoptosis. (Q-V) Transverse sections of zebrafish midbrain and retina labeled with an antibody to the DNA damage marker γ-H2AX (green) and nuclear label DAPI (blue). (Q, S, U) Few γ-H2AX labeled cells are present in wildtype midbrain or retina at 2 dpf (n=3) and 3 dpf (n=3). (R, T, V) In the *rbbp4^Δ4/Δ4^* mutants (n=3) at 2 dpf a significantly greater number of cells with γ-H2AX labeling was present in the brain (W, *p*=0.0008) and retina (X, *p*=0.0032). By 3 dpf (n=3) the number was not significantly different than in wildtype (midbrain W, *p*=0.1211; retina X, *p*=0.0537), likely due to loss of cells by apoptosis. (T) Higher magnification view of nuclear morphology and γ-H2AX labeling in the *rbbp4^Δ4/Δ4^* mutant 2dpf retina. Closed arrow shows pannuclear staining; open arrow shows nuclear ring staining; dotted arrow shows foci pattern of staining. Data represent mean ± s.d. *p* values calculated with one-tailed Student’s *t*-test. Scale bars: 100 μm (I, J, M, N, Q, R, U, V); 10 μm (K, L, S, T).

### Neural precursor apoptosis after Rbbp4 loss is dependent on Tp53

To test whether apoptosis in *rbbp4*^Δ4/Δ4^ neural precursors is activated by the Tp53-dependent DNA damage response we knocked down Tp53 activity using an antisense translation blocking *tp53* morpholino (Fig 5A). Tp53 knock down did not affect development or cell viability in the wildtype midbrain or retina at 2 dpf and 3 dpf, but was able to significantly suppress activated Caspase-3 labeling and apoptosis in the *rbbp4*^Δ4/Δ4^ midbrain and retina at 2 dpf (Fig 5B-5K). At 3 dpf,Tp53 knock down significantly reduced activated Caspase-3 labeling in the *rbbp4*^Δ4/Δ4^ retina but not the midbrain (Fig 5J and 5K), which may be due to a greater decrease in midbrain proliferation compared to the retina. These results are consistent with Rbbp4 preventing Tp53-dependent apoptosis in neural precursors. Tp53 knockdown was able to suppress γ-H2AX labeling in the midbrain and retina of 2 dpf *rbbp4*^Δ4/Δ4^ mutant embryos (Fig 5L-5O, 5T and 5U). By 3 dpf the suppressive effect was no longer significant (Fig 5P-5S, 5T and 5U), possibly due to reduced numbers of viable cells in the proliferative regions of the midbrain and retina. Overall, the nearly complete suppression of activated Caspase-3 and γ-H2AX labeling in 2 dpf *rbbp4*^Δ4/Δ4^ mutant embryos after Tp53 knockdown together indicate Rbbp4 is required to prevent persistent DNA damage and activation of Tp53 dependent programmed cell death in neural precursors.

**Fig 5.**
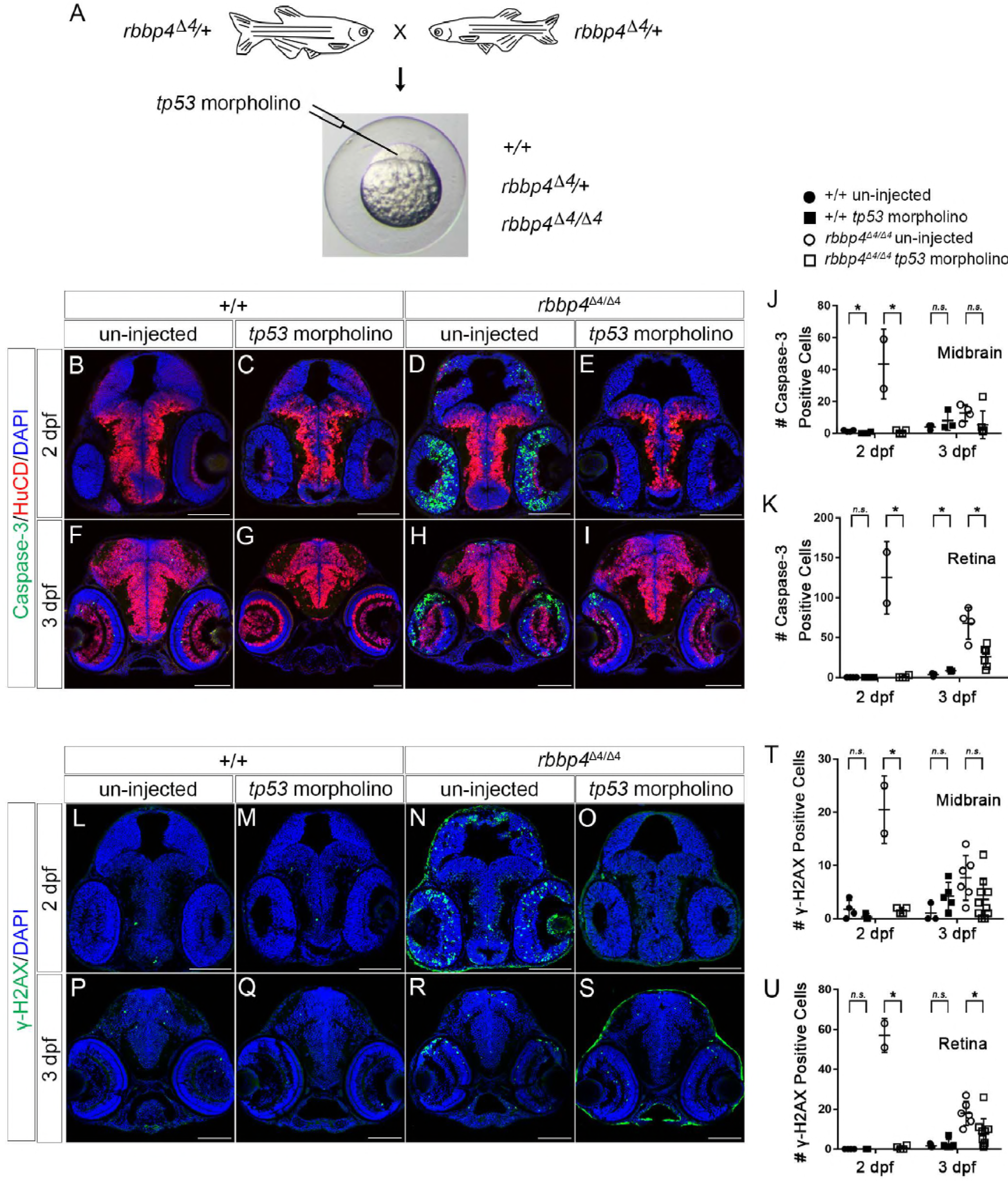
Tp53-dependent activation of DNA damage response and apoptosis in *rbbp4^Δ4/Δ4^* mutant neural precursors. 2 dpf and 3 dpf wildtype and *rbbp4^Δ4/Δ4^* larval midbrain sections after *tp53* knockdown. (A) Inhibition of Tp53 activity by antisense morpholino injection into embryos from *rbbp4^Δ4^/+* adult incross. (B-I) Activated Caspase-3 (green), HuC/D (red) and DAPI (blue) labeling at 2 dpf (B-E) and 3 dpf (F-I). (J) Comparison of Caspase-3 positive cells in the midbrain between un-injected and *tp53* MO injected wildtype at 2 dpf (un-injected n=4; *tp53* MO injected n=4; *p*=0.017), *rbbp4^Δ4/Δ4^* at 2 dpf (un-injected n=2; *tp53* MO injected n=4; *p*=0.0112), wildtype at 3 dpf (un-injected n=3; *tp53* MO injected n=3; *p*=0.3349), and *rbbp4^Δ4/Δ4^* at 3 dpf (un-injected n=4; *tp53* MO injected n=6; *p*=0.1736). (K) Comparison of Caspase-3 positive cells in the retina between un-injected and *tp53* MO injected wildtype at 2 dpf (un-injected n=4; *tp53* MO injected n=4; *p*=0), *rbbp4^Δ4/Δ4^* at 2 dpf (un-injected n=2; *tp53* MO injected n=4; *p*=0.0032), wildtype at 3 dpf (un-injected n=3; *tp53* MO injected n=3; *p*=0.0285), and *rbbp4^Δ4/Δ4^* at 3 dpf (un-injected n=4; *tp53* MO injected n=6; *p*=0.0037). (L-S) γ-H2AX labeling (green), and nuclear stain DAPI (blue) at 2 dpf (K-N) and 3 dpf (O-R). (T) Comparison of γ-H2AX positive cells in the midbrain between un-injected and *tp53* MO injected wildtype at 2 dpf (un-injected n=4; *tp53* MO injected n=3; *p*=0.2344), *rbbp4^Δ4/Δ4^* at 2 dpf (un-injected n=2; *tp53* MO injected n=4; *p*=0.0024), wildtype at 3 dpf (un-injected n=3; *tp53* MO injected n=5; *p*=0.1101), and *rbbp4^Δ4/Δ4^* at 3 dpf (un-injected n=6; *tp53* MO injected n=10; *p*=0.0657). (U) Comparison of γ-H2AX positive cells in the retina between un-injected and *tp53* MO injected wildtype at 2 dpf (un-injected n=4; *tp53* MO injected n=3; p-value=0), *rbbp4^Δ4/Δ4^* at 2 dpf (un-injected n=2; *tp53* MO injected n=4; *p*=0.0001), wildtype at 3 dpf (un-injected n=3; *tp53* MO injected n=5; *p*=0.5784), and *rbbp4^Δ4/Δ4^* at 3 dpf (un-injected n=6; *tp53* MO injected n=10; *p*=0.0128). Data represent mean ± s.d. *p* values calculated with one-tailed Student’s t-test. Scale bars: 100 μm.

### Rbbp4 mutant retinal neural precursors fail to initiate quiescence and differentiation

The results presented above suggest loss of Rbbp4 may induce apoptosis in retinal neural precursors that fail to become quiescent and initiate differentiation. We first used BrdU labeling to follow the fate of *rbbp4^Δ4/Δ4^* mutant neural precursors in the post-embryonic retina. Retinal neurogenesis proceeds in a conveyor belt pattern, with stem cells at the ciliary marginal zone generating progenitors that become progressively more committed as they move inward in the growing retina (28, 29). 2 dpf embryos were exposed to a 3-hour BrdU pulse and collected immediately or chased until 3 dpf and 5 dpf. At the end of 2 dpf in wildtype retina BrdU incorporates into stem cells at the CMZ and progenitors in the inner nuclear and newly developing photoreceptor layers (Fig 6A). A similar pattern of BrdU incorporation was detected in the *rbbp4*^Δ4/Δ4^ mutant retina (Fig 6B). At 3 dpf BrdU labeled cells were found in the region adjacent to the CMZ in both wildtype and *rbbp4*^Δ4/Δ4^ (Fig 6C and 6D outline), however many fewer BrdU positive cells were present in the mutant. By 5 dpf, BrdU positive cells were incorporated into the ganglion and inner nuclear layers of the laminated region of the wildtype retina (Fig 6E outline) but were missing from the *rbbp4*^Δ4/Δ4^ mutant (Fig 6F). In the *rbbp4*^Δ4/Δ4^ mutant, stem cells were still present but lacked BrdU label (Fig 6F arrow). BrdU labeled cells in the central part of the *rbbp4*^Δ4/Δ4^ mutant retina (Fig 6F) may represent rod photoreceptor progenitors or early born Mueller glia in the inner nuclear layer. Together, these results suggest neural progenitors or newborn neurons fail to survive in the absence of Rbbp4.

**Fig 6.**
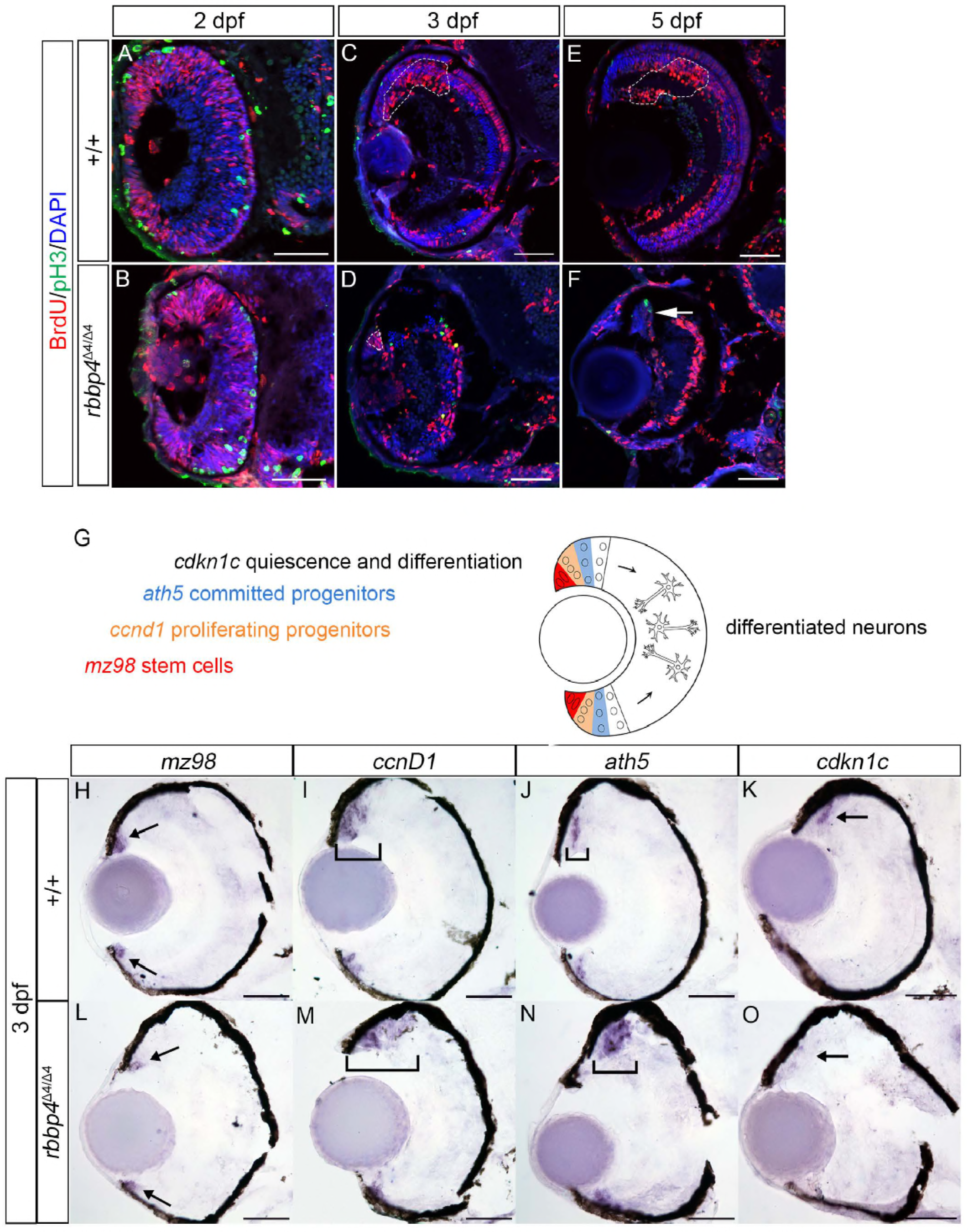
Neural precursors fail to survive and initiate quiescence and differentiation in the *rbbp4^Δ4/Δ4^* homozygous mutant retina. (A-F) BrdU pulse-chase labeling to examine the fate of new born neurons in the *rbbp4^Δ4/Δ4^* mutant retina. (A, B) 2 dpf zebrafish embryos treated with a 2.5 hour BrdU pulse and immediately sacrificed. Transverse sections of retina from wildtype (n=4) and *rbbp4^Δ4/Δ4^* mutants (n=5) were labeled with antibodies to BrdU and mitotic marker phospho-Histone H3 (pH3). BrdU labels proliferating cells at the retina ciliary marginal zone and cells scattered throughout the laminating retina. (C,D) Pulse-chased larvae at 3dpf. In wildtype retina (n=8) BrdU labeled cells are located in the region adjacent to the cmz where neural precursors and newly differentiated neurons reside (C outline). A small section of BrdU labeled cells remains at the ciliary marginal zone in the *rbbp4^Δ4/Δ4^* mutant retina (n=6) (D outline). (E,F) Pulse-chased larvae at 5 dpf. In wildtype (n=7) the BrdU labeled cells are now more centrally located in an older region of the growing retina (E outline). *rbbp4^Δ4/Δ4^* mutant retina (n=7) older born neurons in the central retina maintain BrdU labeling, however, BrdU-labeled newborn neurons are absent. BrdU-negative stem cells persist at the ciliary marginal zone in the *rbbp4^Δ4/Δ4^* mutant (F arrow). (G) Diagram of pattern of gene expression marking stages of neurogenesis in the developing zebrafish retina. (H-O) Transverse sections of 3 dpf zebrafish retina labeled by *in situ* hybridization to examine expression of the stem cell marker *mz98* (H, L arrows), proliferating progenitor marker *ccnD1* (I, M brackets), committed progenitor marker *ath5* (J, N brackets) and differentiating precursor marker *cdkn1c* (K, O arrows). (H-K) Wildtype retina (n=5) shows location of progressively committed precursor cell populations. (L-O) In the *rbbp4^Δ4/Δ4^* homozygous mutant retina (n=3) the proliferating and committed neural progenitor cell populations are expanded compared to wild type. *cdkn1c* expression is absent in the *rbbp4^Δ4/Δ4^* mutant retina (O arrow). Scale bars: 50 μm.

Neurogenesis in the zebrafish retina can be visualized by expression of genes in discreet sectors that mark progressive steps in neural progenitor commitment (30). To identify the stage at which *rbbp4*^Δ4/Δ4^ neural precursors are lost we performed *in situ* hybridization on 3dpf wildtype and *rbbp4*^Δ4/Δ4^ retina with probes to collage type XV alpha 1b (*mz98*) (31), cyclin D1 (*ccnD1*) (32), atonal 7 (*ath5*) (33) and cyclin dependent kinase inhibitor 1Ca (*cdkn1c*) (34). In wildtype, cells at the periphery of the ciliary marginal zone express stem cell marker *mz98* (Fig 6H arrow), followed by proliferating progenitors that express *ccnD1* (Fig 6I bracket), and then *ath5*-expressing committed neural precursors (Fig 6J bracket). Lastly, *cdkn1c* expression marks precursors arrested in G1 that will initiate quiescence and differentiation (Fig 6K arrow). In the *rbbp4*^Δ4/Δ4^ mutant retina *mz98*-expressing cells were present in the CMZ (Fig 6L arrow). The population of *ccnd1*-labeled progenitors and *ath5*-expressing committed precursors was significantly expanded in the mutant compared to wildtype (Fig 6M and 6N brackets). In addition, cdkn1c-expression was not detected in the next adjacent sector (Fig 6O arrow). These results indicate that in the absence of Rbbp4, *ath5*-expressing committed precursor loss occurs before the transition when cells exit the cell cycle to initiate quiescence and differentiation.

### Rbbp4 is required for survival of *rb1-*mutant retinal neural precursors

We previously demonstrated that in *rb1* mutant zebrafish larvae, neural progenitors can re-enter the cell cycle, and phospho-Histone H3 labeled M-phase cells can be detected throughout the *rb1* mutant brain and retina (11). To determine whether survival of *rb1* mutant neural precursors is dependent on Rbbp4, we tested whether *rbbp4*; *rb1* double mutant neural precursors undergo apoptosis (Fig 7A-7H, Fig S7). At 3dpf, in the *rbbp4*^Δ4/Δ4^ mutant retina activated caspase-3 and fragmented nuclei are observed in neural progenitors in the region adjacent to the ciliary marginal zone (Fig 7F dashed outline). Activated caspase-3 labeling is absent from rb1^Δ7/Δ7^ mutant retina and brain, however, in the *rbbp4*^Δ4/Δ4^; rb1^Δ7/Δ7^ double mutant, activated caspase-3 is present, similar to the single *rbbp4*^Δ4/Δ4^ mutant (Fig 7G and 7H). Labeling with phospho-Histone H3 antibodies confirm the rb1^Δ7/Δ7^ single and rbbp4^Δ4/Δ4^; rb1^Δ7/Δ7^ double mutants have increased numbers of cells in mitosis compared to wildtype brain and retina (Fig 7I-7P). These results show that loss of Rbbp4 can induce apoptosis in *rb1* mutant neural precursors, indicating the viability of *rb1* mutant cells is dependent Rbbp4 (Fig 7Q).

**Fig 7.**
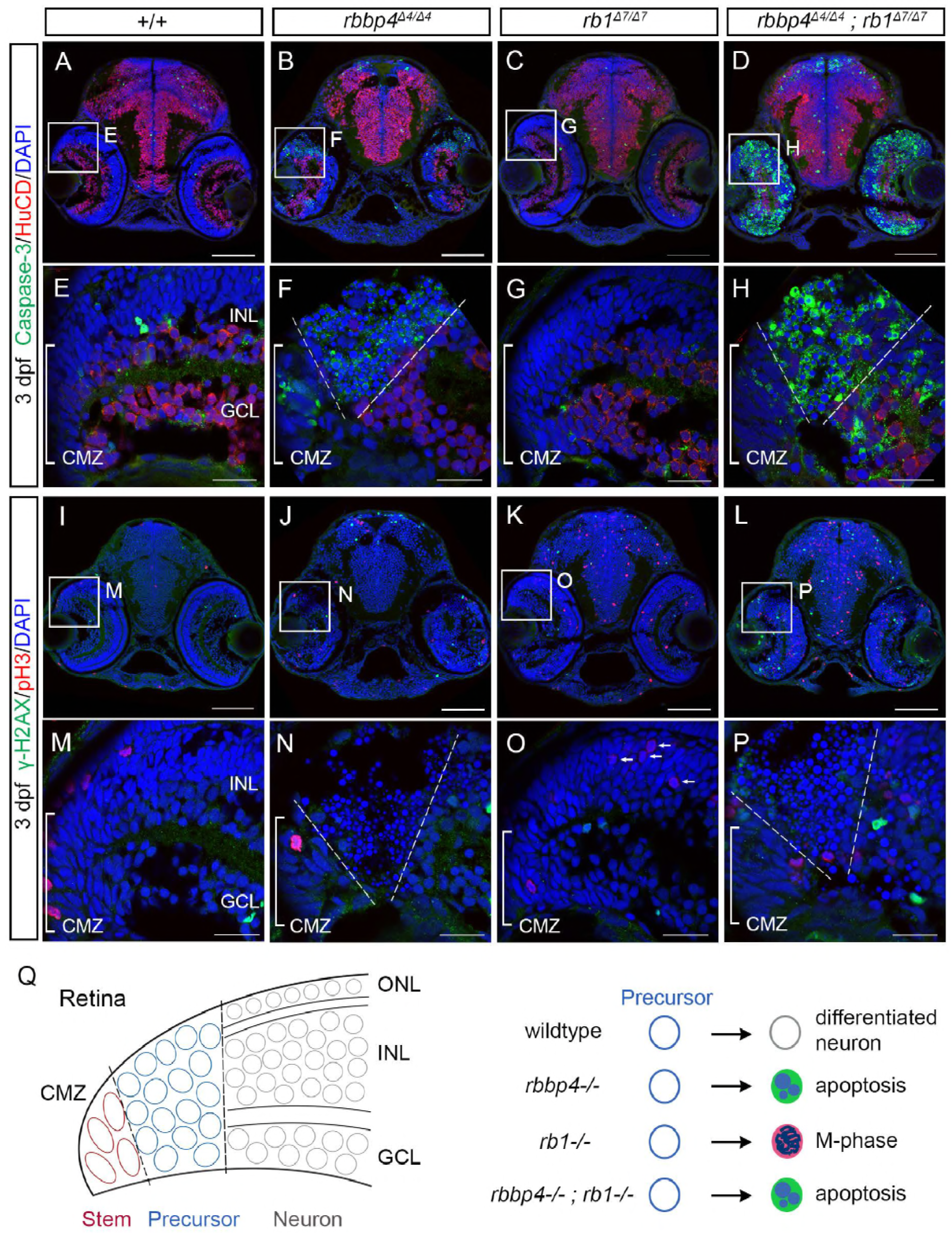
Rbbp4 is required for survival of *rb1/rb1* mutant retinal neural precursors. 3 dpf wild type +/+ (n=3), *rbbp4^Δ4/Δ4^* (n=3), rb1^A7/A7^ (n=4), and rbbp4^Δ4/Δ4^; rb1^A7/A7^ (n=2) siblings from a rbbp4^Δ4/+^; rb1^Δ7/+^ incross were sectioned and labeled with antibodies to activated caspase-3 and HuCID (A-H) or γ-H2AX and _hosphor-Histone H3 (I-P). Activated caspase-3 and nuclear fragmentation are present in the region containing neural precursors (dashed outline) adjacent to the ciliary marginal zone in both *rbbp4^Δ4/Δ4^* and rbbp4^Δ4/Δ4^; rb1^Δ7/Δ7^ mutants. (Q) Diagram modeling requirement for Rbbp4 and Rbl in retinal neurogenesis. Newborn retinal neural precursors adjacent to the ciliary marginal zone will differentiate into neural cell types that populate the retina ganglion cell, inner nuclear, and outer nuclear layers. Both Rbbp4 and Rb1 are required for neural precursors to initiate quiescence and differentiation, however, loss of *rbbp4* leads to DNA damage-induced apoptosis, while loss of *rb1* allows entry into mitosis. Double mutant precursors undergo apoptosis, indicating *rb1* mutant precursor viability is dependent on Rbbp4. CMZ ciliary marginal zone; GCL ganglion cell layer; INL inner nuclear layer; ONL outer nuclear layer. Scale bars: A-D, I-L 100 μm; E-H, M-P 20 μm.

## Discussion

In this study we demonstrate *in vivo* Rbbp4 is essential for vertebrate neurogenesis and is required for neural stem cell proliferation and neural precursor survival. While Rbbp4 was originally identified as a binding partner of the Rb1 tumor suppressor, we show for the first time that survival of *rb1*-mutant neural precursors is dependent on *rbbp4*. We find that Rbbp4 is overexpressed in *rb1*-embryonal brain tumors and is upregulated across the spectrum of human embryonal and glial tumor types. Together with recent studies examining the role of Rbbp4 in glioblastoma DNA damage repair (9) and neuroblastoma tumor progression (10), our study provides additional evidence that Rbbp4 might contribute to brain tumor cell survival by promoting proliferation and circumventing DNA damage induced apoptosis.

Our analysis of zebrafish *rbbp4* mutants revealed Rbbp4 is necessary for neurogenesis with distinct roles in neural stem and progenitor cells. Loss of Rbbp4 did not affect stem cell survival, however, BrdU lineage tracing showed mutant stem cells underwent a limited number of divisions before exiting the cell cycle. The morphology of the stem cells was abnormally enlarged, which could represent the onset of senescence that is characterized by cellular hypertrophy (35). These results are consistent with other *in vivo* and *in vitro* studies indicating a role for Rbbp4 in preventing cellular senescence. Rbbp4 is a component of the DREAM/MuvB complex that controls quiescence by promoting cell cycle gene expression and progression through G2 (36), but has also been shown to be required for oncogene-induced senescence in immortalized BJ fibroblasts (37). In Arabidopsis, the Rbbp4 homolog *AtMSI1* is necessary for persistent growth of reproductive tissues through expression of homeotic genes and heterochromatin assembly (38). In addition to controlling senescence through chromatin structure and gene expression, Rbbp4 has been implicated in blocking cellular senescence through the regulation of nuclear import (39). Lastly, cancer cells that maintain telomere length using the alternative lengthening of telomere (ALT) pathway recruit NuRD to their telomeres through the transcription factor ZNF827 (40). Recently it was shown that Rbbp4 binds ZNF827 (41), suggesting Rbbp4 might contribute to preventing cancer cell replicative senescence by association of NuRD with telomeres. Together these studies suggest a critical role for Rbbp4 in stem cell maintenance, possibly through multiple nuclear processes, including transcriptional regulation of cell cycle progression, heterochromatin formation, telomere lengthening, and nuclear transport.

A major finding in our study is the *in vivo* requirement for Rbbp4 to prevent Tp53-dependent apoptosis in zebrafish neural progenitors in the growing midbrain and retina. In the *rbbp4* mutant retina, the expansion of the *ath5-*positive progenitor population and the absence of *cdkn1c*-expressing cells suggests progenitors fail to transition to a neural precursor and initiate quiescence. The accumulation of γ-H2AX labeling in *rbbp4* mutant progenitors would suggest persistent DNA damage activates the DNA damage response and Tp53-dependent apoptosis. Given the role of Rbbp4 in multiple chromatin remodelers that regulate gene expression, the primary effect of loss of Rbbp4 may be altered transcriptional regulation. This could affect multiple pathways including the DNA damage response, as has been shown in glioblastoma multiforme cells (9), leading to persistent DNA damage and activation of Tp53 apoptosis. Alternatively, loss of Rbbp4 could directly affect genome integrity in neural precursors. Knockdown of Rbbp4 in cell culture leads to defects in DNA synthesis, nucleosome assembly and heterochromatin formation (42, 43). Similar mechanisms may cause DNA damage in zebrafish *rbbp4* mutant embryos, possibly through disruption of the CAF-1 nucleosome assembly factor that contains p150, p60, and Rbbp4 (44). In addition to the classical γ-H2AX foci representing DNA double strand breaks (27), the majority of γ-H2AX labeling in *rbbp4* mutant embryos appeared as pan-nuclear or nuclear rings, which mark extensive DNA damage and disruption of chromatin integrity (45). γ-H2AX nuclear ring labeling has been described in response to activation of death receptors by TRAIL ligand, indicating extrinsic signaling independent of Tp53 could also contribute to apoptosis in *rbbp4* mutant cells (46). However, our ability to suppress apoptosis after Tp53 knockdown in *rbbp4* mutant embryos would argue that intrinsic activation of Tp53 and the DNA damage response is responsible for apoptosis.

In conclusion, our study indicates Rbbp4 may promote proliferation and survival of wildtype and *rb1* mutant neural precursors by circumventing DNA damage induced apoptosis. Disruption of transcriptional regulation, chromatin assembly and DNA repair could each contribute to apoptosis after loss of Rbbp4. Knockdown of Tp53 blocked apoptosis, providing a means to prevent cell death and examine how loss of Rbbp4 changes the epigenetic landscape and gene expression profile in *rbbp4* mutant neural precursors. These future studies would provide new insight into the mechanism by which Rbbp4 and its associated chromatin remodelers promote proliferation and survival during neurogenesis.

## Materials and Methods

### Zebrafish care and husbandry

Zebrafish were reared in an Aquatic Habitat system (Aquatic Ecosystems, Inc., Apopka, FL). Fish were maintained on a 14-hr light/dark cycle at 27°C. Transgenic lines were established in a WIK wild type strain obtained from the Zebrafish International Research Center (http://zebrafish.org/zirc/home/guide.php). The zebrafish *rbbp4* and *rb1* alleles used in this study were isolated by CRISPR-Cas9 or TALEN targeting and described previously: *rbbp4* 4 base pair (bp) deletion allele *rbbp4Δ4^is60^* (11); *rb1* 7bp deletion allele *rb1Δ7^is54^* (18). For *in situ* hybridization and immunohistochemistry experiments, embryos were collected and maintained at 28.5°C in fish water (60.5 mg ocean salts/l) until harvesting. Embryos were staged according to published guidelines (47). All experimental protocols were approved by the Iowa State University Institutional Animal Care and Use Committee (Log # 11-06-6252-I) and are in compliance with American Veterinary Medical Association and the National Institutes of Health guidelines for the humane use of laboratory animals in research.

### Larval genotyping and *tp53* morpholino injections

Primers to amplify *rbbp4* exon 2 containing the *rbbp4Δ4^is60^* mutation: Forward 5' GCGTGATGACAGATCTCATATTGTTTTCCC 3'; Reverse 5' CTGGTGACATCTGGCAACCACT 3'. Primers to amplify *rb1* exon 2 containing the *rb1Δ7^is54^* mutation: Forward 5’-TTTCCAGACACAAGGACAAGGATCC-3’; Reverse 5’-GCAGATATCAGAAGAAAGAGTACATTTGTCTT-3’. To inhibit Tp53-dependent apoptosis, embryos were injected at the one cell stage with 2 ng of *tp53* translation blocking morpholino GCGCCATTGCTTTGCAAGAATTG (48) (Gene Tools; ZDB-MRPHLNO-070126-7.

### Isolation of transgenic line *Tg*(*Tol2<ubi:rbbp4-2AGFP>*)*^is61^*

The *rbbp4* 1275 bp cDNA minus the translation termination codon was amplified by reverse transcription PCR with Forward 5’-catgTCTAGATGTGGAGTCGTTATGGCTG-3’ and Reverse 5’-catgGGATCCTCCCTGAACCTCAGTGTCTG-3’ primers that included 5’ *Xba*I and 3’ *BamH*I sites, respectively. The *Tol2<ubi:rbbp4-2AGFP>* transgene was assembled using the NEBuilder HiFi DNA Assembly protocol and mix (New England Biolabs E2621S) in the mini-*pTol2* vector (49). The zebrafish *ubiquitin* promoter (50) was cloned into the vector followed by the *rbbp4* cDNA, an in-frame 2A viral peptide GFP cassette (51) and the zebrafish β*-actin* 3’UTR (Dr. Darius Balciunas, Temple University; (52)). 1μg of linearized *pT3TS-Tol2* transposase plasmid (49) was used as template for *in vitro* synthesis of *Tol2* transposase capped, polyadenylated mRNA with T3 mMessage mMachine High Yield Capped RNA transcription kit (Ambion AM1348). Synthesized mRNA was precipitated with LiCl and resuspended in RNase, DNase-free molecular grade water. The *Tg*(*Tol2<ubi:rbbp4-2AGFP>*)*^is61^* transgenic line was isolated by co-injection into 1-cell WIK zebrafish embryos of 25pg *Tol2* vector plus 100pg *Tol2* mRNA. 14 founder fish were screened for germline transmission of ubiquitously expressed GFP. A single F1 adult that showed Mendelian transmission of the *Tol2<ubi:rbbp4-2AGFP>* transgene to the F2 generation was used to establish the line.

### Immunohistochemistry

Embryonic (2 dpf) or larval (3-5 dpf) zebrafish were anesthetized in MS-222 Tricaine Methanesulonate and head and trunk dissected. Trunk tissue was placed in 20μl 50mM NaOH for genotyping. Heads were fixed in 4% paraformaldehyde overnight at 4°C, incubated in 30% sucrose overnight at 4°C, then processed and embedded in Tissue-Tek OCT (Fisher 4583). Processing and embedding of zebrafish wildtype adult brain and *rb1-*brain tumor tissue was as described (18). Immunolabeling was performed on TALEN targeted *rb1-*brain tumors from three fish; tumor tissue from two of these fish (individuals #7 and #10) was also used for RNA-Seq libraries in our previous analysis of the *rb1-*embryonal brain tumor transcriptome (11). Tissues were sectioned at 14–16 μm on a Microm HM 550 cryostat. For BrdU labeling experiments, 2 dpf embryos were incubated in 5 μM BrdU (Sigma B5002) in embryo media (53) for 2.5 hours, placed in fresh fish water, then sacrificed immediately or at 3 dpf and 5 dpf. To aid BrdU antigen retrieval tissues were pretreated with 2 M HCl. For RBBP4 labeling, antigen retrieval was performed by placing slides in boiling 10 mM NaCitrate and allowing the solution to cool to room temperature. Antibodies used for labeling: rabbit polyclonal anti-RB Binding Protein 4 RBBP4 1:200 (Bethyl A301-206A-T, RRID: AB_890631); rabbit polyclonal anti-SOX2 1:200 (EMD Millipore AB5603, RRID:AB_2286686); mouse monoclonal anti-proliferating cell nuclear antigen PCNA 1:300 (Sigma P8825); mouse monoclonal anti-glial fibrillary acid protein GFAP 1:1000 (Zebrafish International Research Center zrf-1); rabbit polyclonal anti-phospho-Histone H3 PH3 1:1000 (Cell Signaling Technology; 9701); mouse monoclonal anti-phospho-Histone H3 (Ser10), clone 3H10 1:500 (Millipore 05-806); rabbit polyclonal anti-Brain Lipid Binding Protein BLBP 1:200 (Abcam ab32423); mouse monoclonal anti-HuC/D 1:500 (Invitrogen A-21271); rabbit polyclonal anti-gamma H2A histone family, member X γ-H2AX 1:200 (GeneTex GTX127342); rabbit polyclonal anti-CASPASE-3 1:500 (BD Biosciences 559565); mouse monoclonal anti-SV2 1:100 (Developmental Studies Hybridoma Bank AB_2315387); rabbit polyclonal anti-CALRETININ 1:1000 (Millipore AB5054); mouse monoclonal anti-BrdU 1:500 (Bio-Rad MCA2483); Alexa-594 (Invitrogen A-11005) and Alexa-488 (Invitrogen A-11008) conjugated secondary antibodies 1:500. Tissues were counterstained with 5 μg/ml DAPI and mounted in Fluoro-Gel II containing DAPI (Electron Microscopy Sciences 17985-50) and imaged on a Zeiss LSM700 laser scanning confocal.

### *in situ* hybridization and alcian blue staining

cDNA was amplified by reverse transcription-polymerase chain reaction out of total RNA isolated from wild-type 5 day post-fertilization embryos and cloned into the pCR4-TOPO vector (Invitrogen). Primers for amplification: *mz98* forward 5'-CCGGACACTACACACTCAATGC-3', *mz98* reverse 5'-GTGCTGGATGTAGCTGTTCTCG-3'; *ccnD1* forward 5'- GCGAAGTGGATACCATAAGAAGAGC-3', *ccnD1* reverse 5'- GCTCT GATGTATAGGCAGTTTGG-3'; *ath5* forward 5'-GATTCCAGAGACCCGGAGAAG-3', *ath5* reverse 5'-CAGAGGCTTTCGTAGTGGTAGGAG-3'; *cdkn1c* forward 5'- CGTGGACGTATCAAGCAATCTGG-3', *cdkn1c* reverse 5’-GTCTGTAATTTGCGGCGTGC-3'. Digoxigenin-labeled probes were synthesized using T3 RNA polymerase (Roche #11031163001) and DIG RNA labeling mix (Roche #11277073910) according to the manufacturer’s instructions and stored in 50% formamide at-20°C. Embryonic and larval zebrafish tissues were fixed in 4% paraformaldehyde and embedded in Tissue-Tek OCT (Fisher 4583). *in situ* hybridization was performed on 14–16 μm cryosections. For alcian blue staining of cartilage, zebrafish larvae were anesthetized and fixed in 4% paraformaldehyde overnight at 4°C and incubated in 0.1% alcian blue solution overnight at 4°C. Embryos were rinsed in acidic ethanol and stored in 70% glycerol. Whole larvae and tissue sections were imaged on a Zeiss Discovery.V12 stereomicroscope or Zeiss Axioskop 2 microscope and photographed with a Nikon Rebel camera.

### Western blotting

Whole protein extract was generated by dounce homogenization of ~70 5 dpf embryos in 200 uL Cell Extraction Buffer (ThermoFisher FNN0011). Lysates were run on a NuPAGE™ 12% Bis-Tris Protein Gel (Invitrogen NP0342BOX) and blotted onto PVDF membrane (BioRAD 1620177). Membranes were incubated in either rabbit polyclonal anti-RB Binding Protein 4 RBBP4 (Bethyl A301-206A-T) at 1:800 or mouse monoclonal anti-β-actin (Sigma A2228) at 1:5000 overnight at 4°C, followed by Anti-Rabbit IgG F(ab')2-Alkaline Phosphatase (Sigma SAB3700833) or Anti-Mouse IgG F(ab')2-Alkaline Phosphatase (Sigma SAB3700994) at 1:500 for one hour at room temperature. Membranes were developed with CSPD (Roche 11655884001) and imaged on a BioRAD Molecular Imager ChemiDoc XRS System.

### Quantification and Statistical Analysis

Statistical analysis was performed using GraphPad Prism software. Data plots represent mean ± s.d. *ρ* values were calculated with one-tailed Student’s *t*-test. Statistical parameters are included in the Figure legends.

## Acknowledgements

This authors which to thank Trevor Weiss, Daniel Splittstoesser and Kristen Wall for technical assistance.

## Author Contributions

L.E.S. and M.M. conceived and designed the experiments and wrote the manuscript. L.E.S., M.P.A. and W.A.W. built reagents, performed and documented experiments, and analyzed results. M.K. performed human tumor expression analysis.

## Supporting Information

**S1 Fig.**
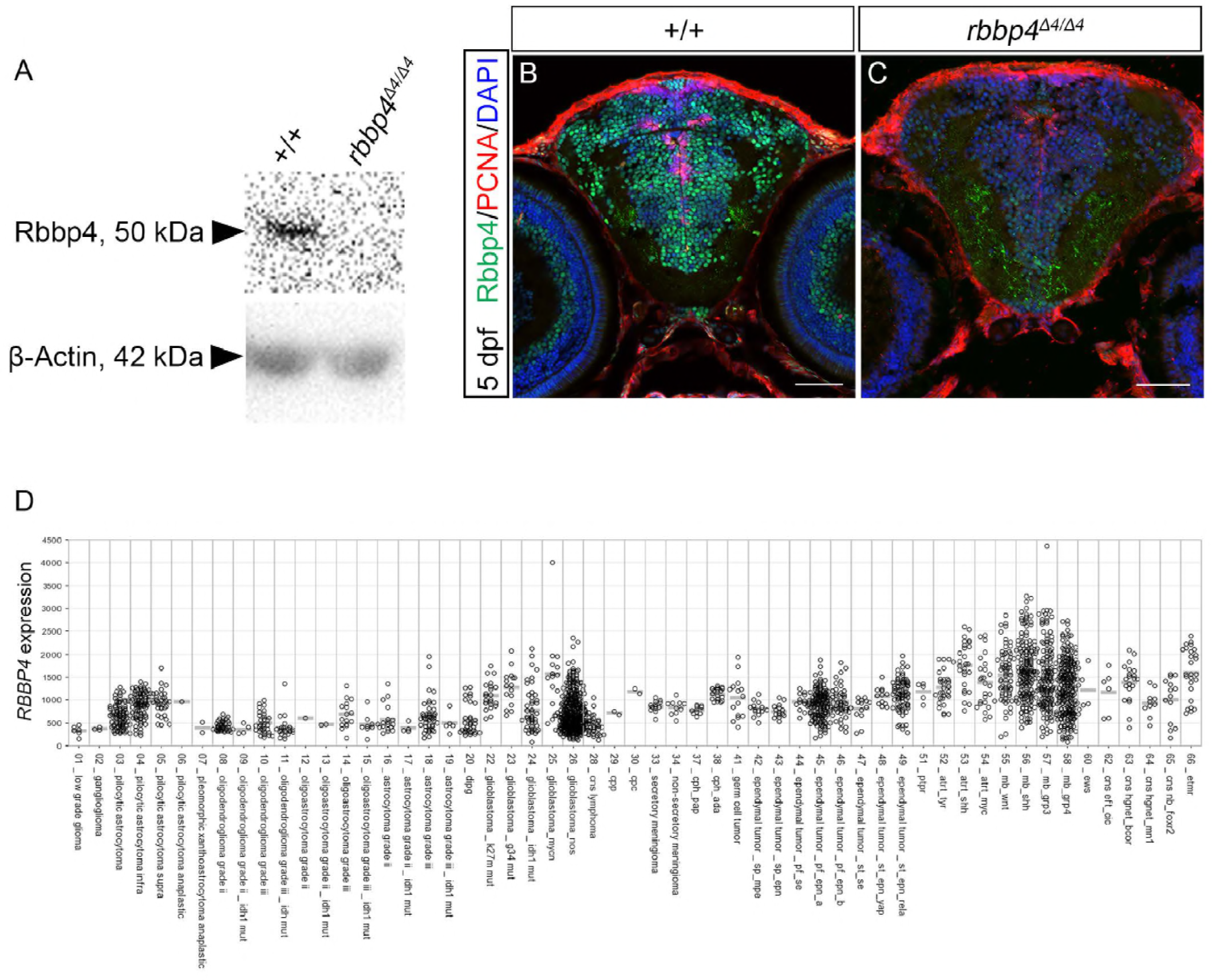
Related to Figs 1 and 2. (A-C) Rbbp4 protein is absent in *rbbp4^Δ4/Δ4^* homozygous mutant larvae at 5 dpf. (D) *RBBP4* is highly expressed in many human brain tumor entities. (A) Western blot of whole protein lysate from wildtype +/+ and homozygous mutant *rbbp4^Δ4/Δ4^* 5 dpf larvae probed with a rabbit polyclonal anti-Rbbp4 antibody. The antibody recognizes a single polypeptide band of ~50kDA, close to the Rbbp4 predicted size of 48kDa. The band is absent from the *rbbp4Δ4/Δ4* mutant protein extract. Lower panel, loading control blot probed with mouse monoclonal antibody β-actin. (B, C) Rbbp4 and proliferative marker PCNA antibody labeling of transverse sections through the head of 5 dpf wildtype (B) and *rbbp4Δ4/Δ4* homozygous mutant (C) zebrafish larvae. Confocal images for each larva were captured at approximately the same gain settings, with a slightly higher gain setting for the homozygote, to confirm the absence of Rbbp4 signal. In wildtype brain (B) high levels of Rbbp4 are present in the nucleus of neurons in the tectum and thalamic regions, but are nearly absent in the *rbbp4Δ4/Δ4* homozygote (C). Scale bars: 50 μm. (D) *RBBP4* Affymetrix data from 2284 human tumor samples (German Cancer Research Center DKFZ). atrt atypical teratoidIrhabdoid tumor, cns central nervous system; cpc choroid plexus carcinoma; cph, cpp choroid plexus papilloma; dipg diffuse intrinsic pontine glioma; etmr embryonal tumor with multilayered rosettes; ews Ewing’s sarcoma; mb medulloblastoma; ptpr papillary tumor of the pineal.

**S2 Fig.**
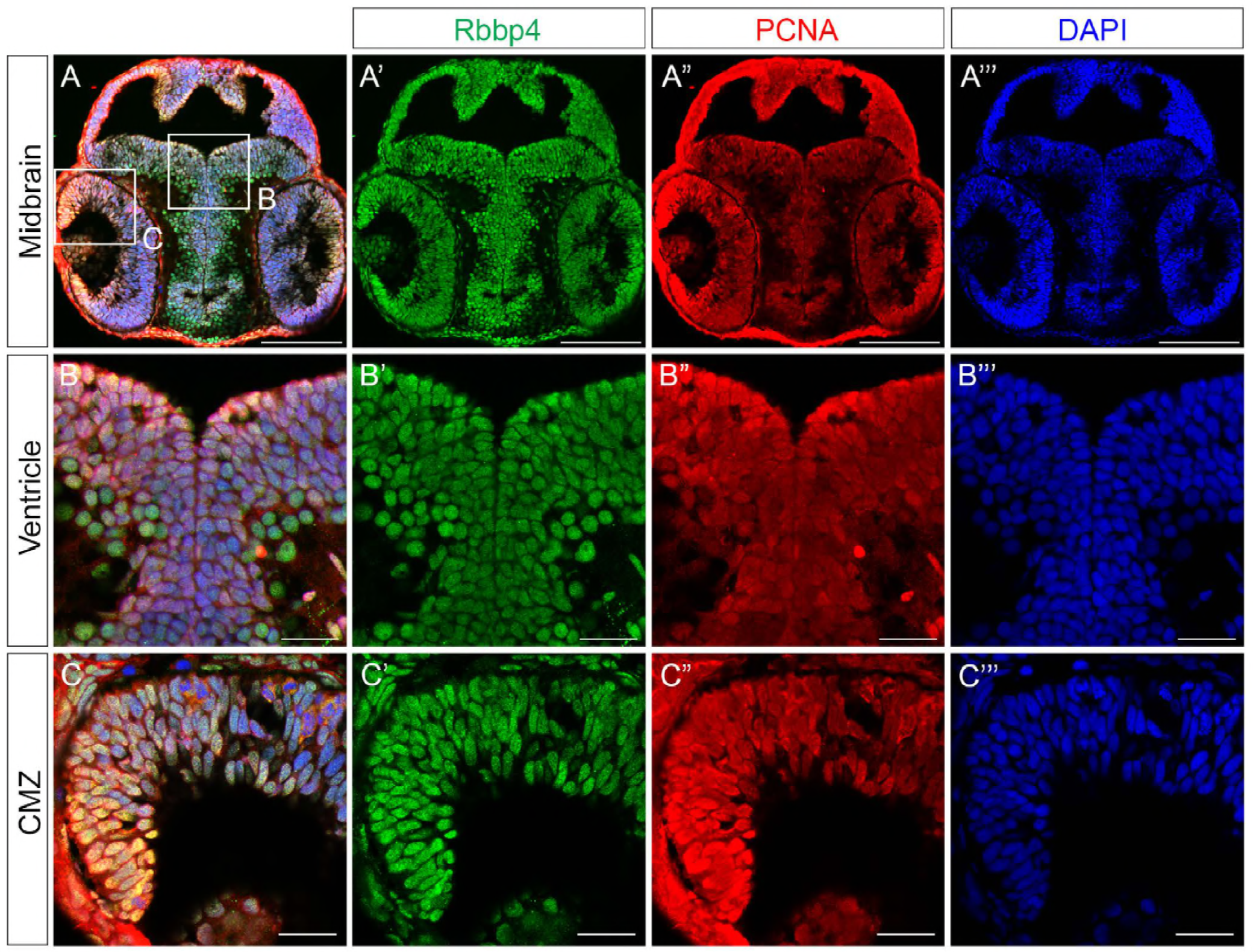
Related to Fig 2. Rbbp4 is ubiquitously expressed in neuroblasts in the embryonic zebrafish brain and retina. (A-A’’’) Representative transverse cryosections (n=3) through 2 dpf wildtype zebrafish midbrain labeled with antibodies to Rbbp4 (green), proliferation maker PCNA (red), and nuclear stain DAPI (blue). Boxes outline the ventricle at the top of the thalamus in the midbrain (B) and the ciliary marginal zone (CMZ) of the retina (C). (B-B’’’) Higher magnification view of the midbrain ventricle at the thalamus. (C-C’’’) Higher magnification view of the ciliary marginal zone of the retina (C-C’’’). Scale bars: 100 μm A-A’’’; 20 μm B-C’’’.

**S3 Fig.**
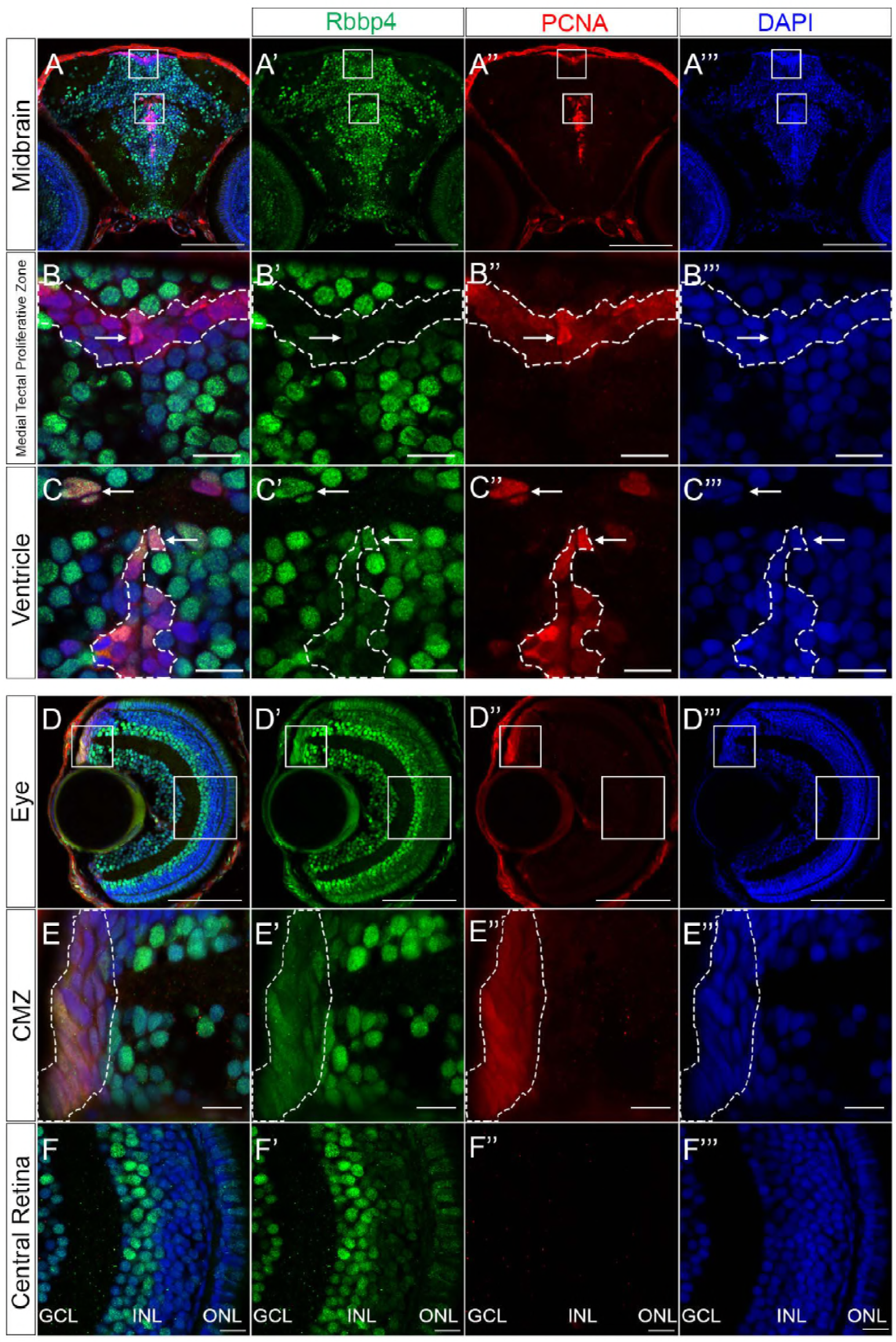
Related to Fig 2. Rbbp4 levels vary in neural stem cells, precursors and neurons in the larval zebrafish brain and retina. Representative transverse cryosections (n=3) of 5 dpf wildtype zebrafish midbrain (A-A’’’) and retina (D-D’’’) labeled with antibodies to Rbbp4 (green), proliferation maker PCNA (red), nuclear stain DAPI (blue). Higher magnification view of the medial tectal proliferative zone (B-B’’’), brain ventricle (C-C’’’), ciliary marginal zone (E-E’’’), and central laminated layers of the retina (F-F’’’). The highly proliferative, PCNA+ population of cells are outlined; arrows indicate cells co-labeled with Rbbp4 and PCNA. Rbbp4 is enriched in a subset of neurons in the midbrain and vitreal side of the retinal inner nuclear layer. GCL, ganglion cell layer; INL, inner nuclear layer; ONL, outer nuclear layer. Scale bars: 100 μm A-A’’’, D-D’’’; 10 μm B-B’’’, C-C’’’, E-E’’’, F-F’’’.

**S4 Fig.**
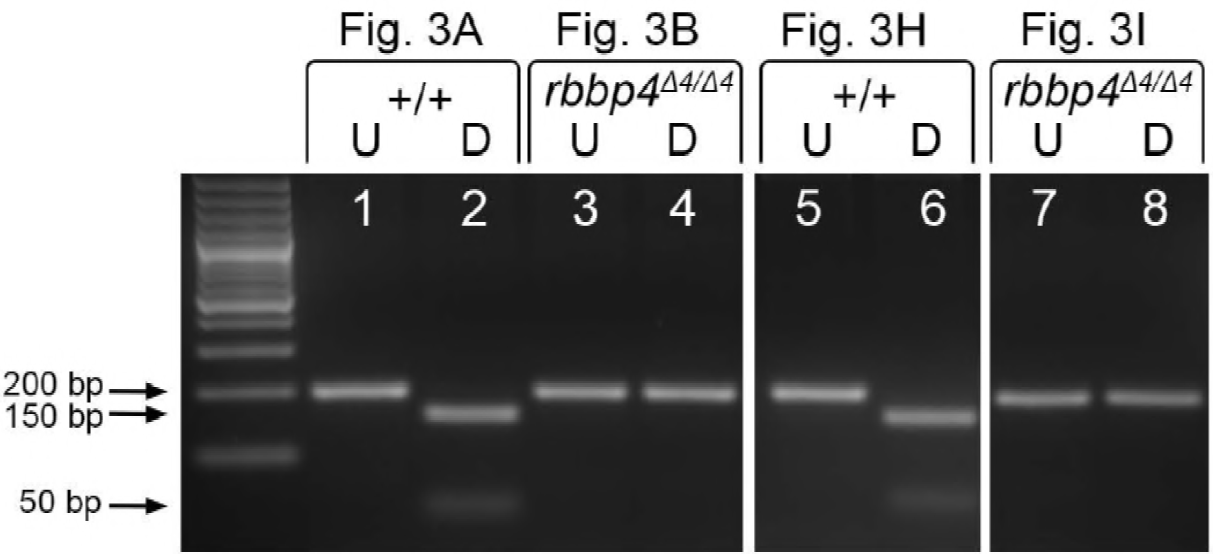
Related to Fig 3. Genotype confirmation of progeny from *rbbp4Δ4/Δ4*; *Tg*(*Tol2<ubi:rbbp4-2AGFP>*) rescue experiment. A 196 bp PCR amplicon surrounding *rbbp4* exon 2 containing the 4 bp deletion mutation that overlaps a *SmlI* restriction enzyme site. (Lanes 1, 2, 5, 6) *SmlI* digestion of +/+ wildtype larva (Fig 3A) and transgenic +/+ larva (Fig 3H) generates 150 and 46 bp fragments. (Lanes 3, 4) In homozygous *rbbp4Δ4/Δ4* larvae (Fig 3B) the *rbbp4Δ4* allele is resistant to digestion. (Lanes 7, 8) The resistant band in lane 8 confirms the *Tg*(*Tol2<ubi:rbbp4-2AGFP>*) rescued larva (Fig 3I) is homozygous for the *rbbp4Δ4* allele. U, un-digested; D, SmlI digested.

**S5 Fig.**
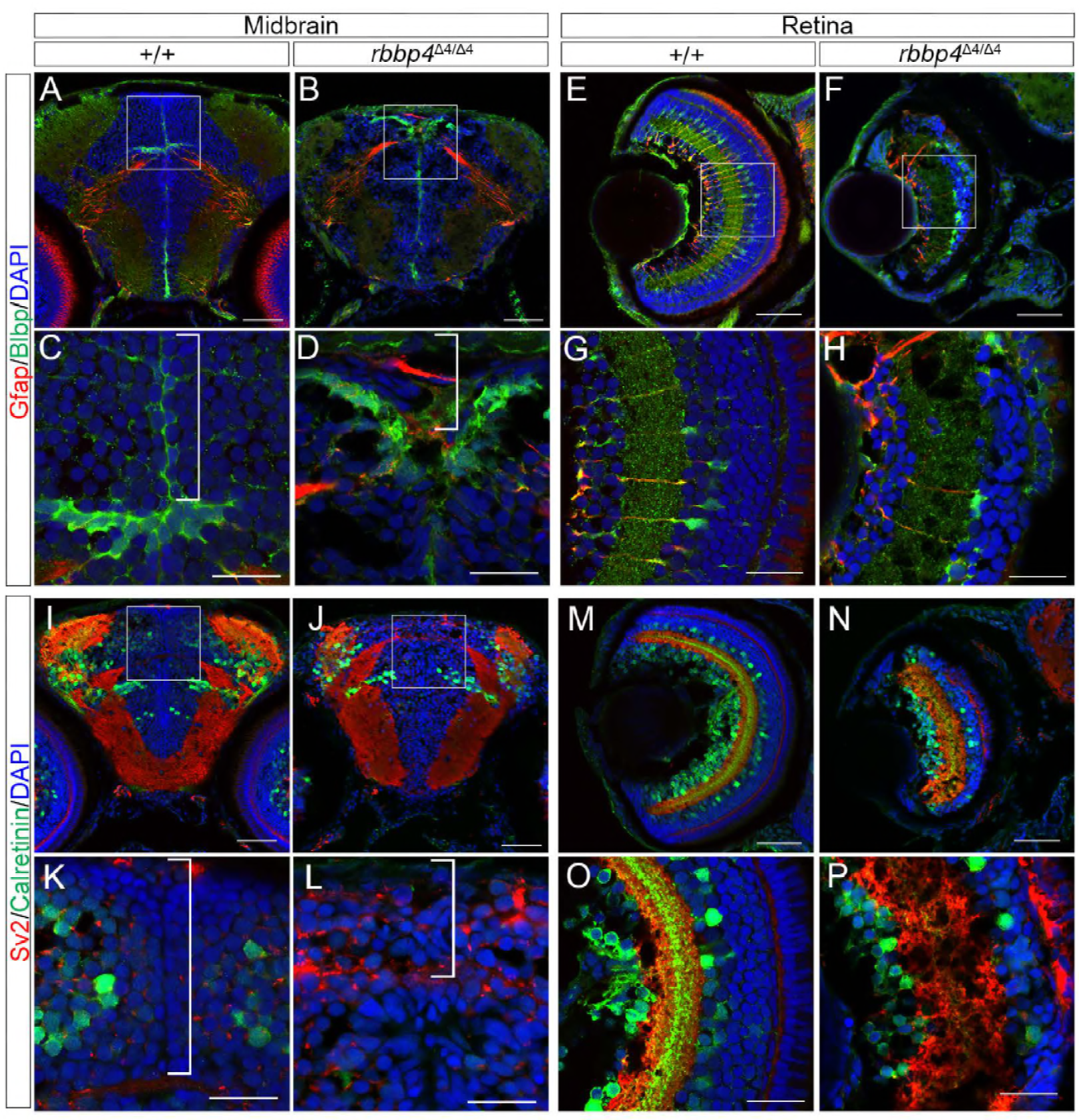
Related to Fig 3. Maternal Rbbp4 allows early born neurons and glia in *rbbp4Δ4/Δ4* homozygotes to undergo normal neurogenesis and differentiation. Transverse section of midbrain and retina from wild type +/+ (n=3) and *rbbp4Δ4/Δ4* homozygous (n=3) 5 dpf larva labeled with antibodies to glial markers Gfap (red) and Blbp (green) and nuclear stain DAPI (blue). (A, C) In wild type midbrain Blbp and Gfap label radial glia cell bodies and projections in the tectum (C bracket) and thalamic regions, respectively. (B, D) The *rbbp4Δ4/Δ4* mutant midbrain contains mature radial glia in the thalamic region (B). Intense Blbp labeling is present where hypertrophic stem cells are located at the ventricle (D bracket). (E, G) Blbp and Gfap label Mueller glia cells bodies and projections across the inner plexiform layer of the wild type retina. (F, H) Blbp and Gfap label persists in early born glia in the *rbbp4Δ4/Δ4* mutant retina. (I-P) Transverse section of midbrain and retina from wild type +/+ (n=3) and *rbbp4Δ4/Δ4* mutant (n=3) 5 dpf larvae labeled with antibodies to the synaptic vesicle marker Sv2 (red) and interneuron and ganglion cell marker Calretinin (green) and nuclear stain DAPI (blue). (I, K) In wildtype brain, Sv2 labels the neuropil, and Calretinin labels the cell bodies of a subset of mature neurons in the thalamus and tectum (K bracket). (J, L) The *rbbp4Δ4/Δ4* mutant shows a reduction in the tectal neuropil and number of Calretinin-positive neurons (L bracket). (M, O) Sv2 and Calretinin labeling in wild type retina. (N, P) Early born neurons maintain Calretinin labeling and Sv2 labeled projections across the inner and outer plexiform layers in the *rbbp4^Δ4/Δ4^* mutant retina. Scale bars: 50 μm A, B, E, F, I, J, M, N; 20 μm C, D, G, H, K, L, O, P.

**S6 Fig.**
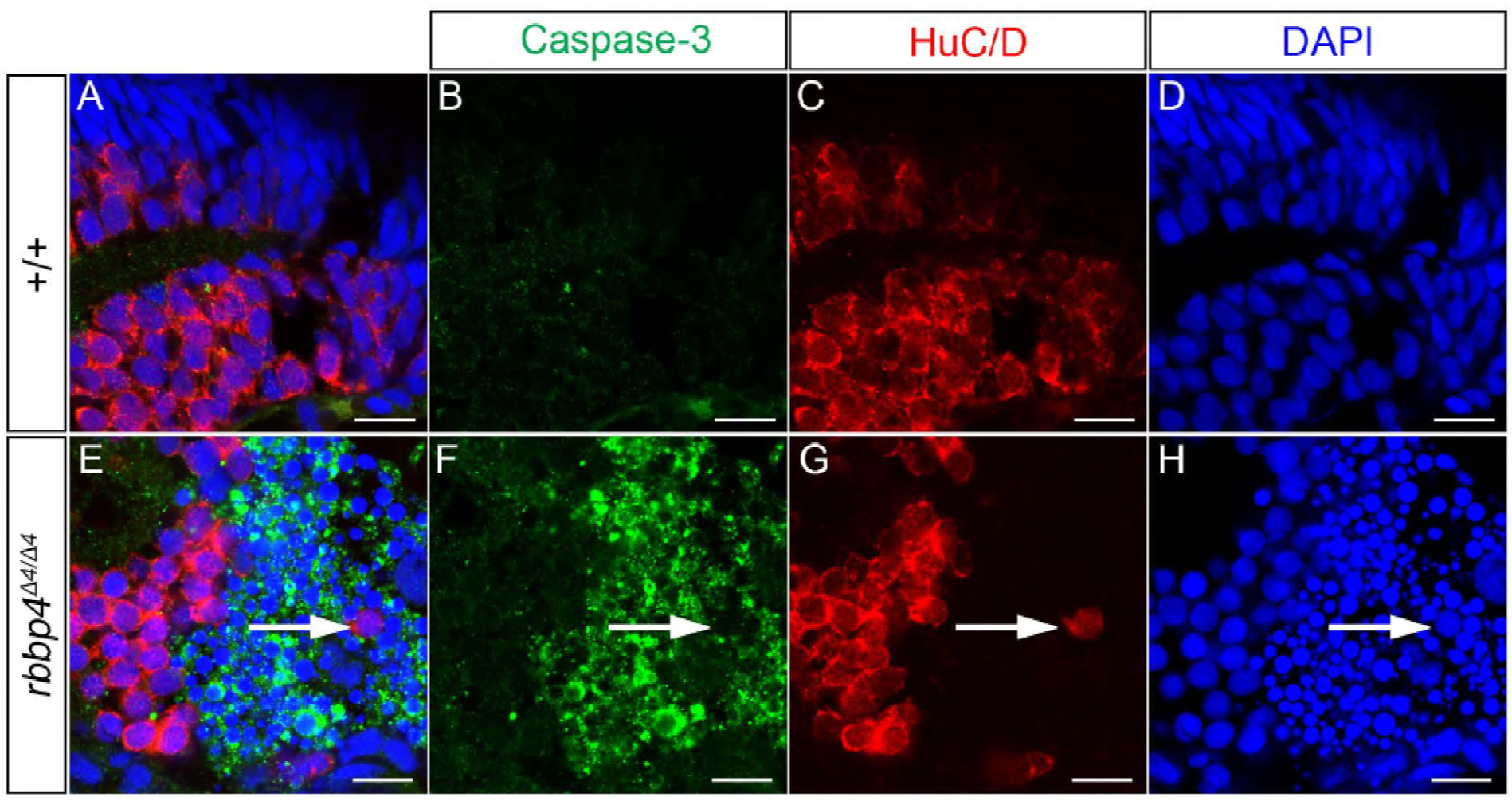
Related to Fig 4. Caspase-3 labeled rbbp4*^Δ4/Δ4^* mutant retinal cells undergoing apoptosis do not express the neuronal marker HuC/D. (A-H) Transverse sections of 3 dpf zebrafish retinas labeled with antibodies to the apoptosis marker activated Caspase-3 (green), neuronal marker HuC/D (red), and nuclear stain DAPI (blue). (A-D) In wildtype +/+ HuC/D is detected in neuron cell bodies in the inner nuclear layer and photoreceptor layer of the retina. (E-H) In the *rbbp4Δ4/Δ4* homozygous retina cells expressing activated Caspase-3 do not co-label with HuC/D. Arrow points to an activated Caspase-3-negative/HuC/D-positive cell surrounded by activated Caspase-3-positive/HuC/ID-negative cells. Scale bars: 100 μm.

**S7 Fig.**
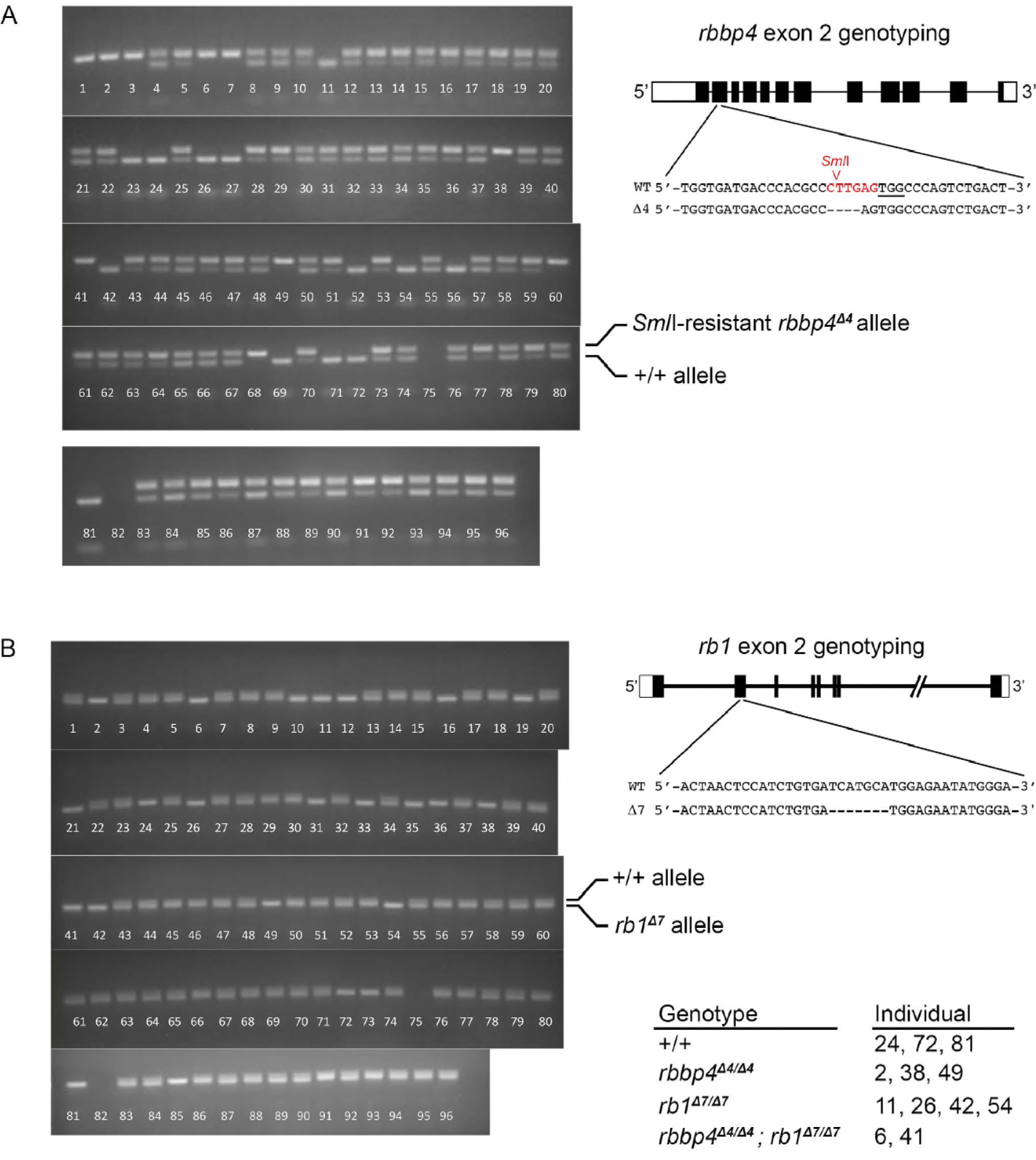
Related to Fig 7. PCR genotyping to confirm genotype of +/+ wildtype, *rbbp4Δ4/Δ4* mutant, *rb1Δ7*/*Δ7* mutant, and *rbbp4Δ4/Δ4; rb1Δ7*/*Δ7* double mutant 3 dpf larvae from *rbbp4Δ4/+; rb1Δ7/+* double heterozygous incross. Each lane corresponds to an individual progeny. A Gels of rbbp4 exon 2 PCR amplicons digested with *Sml*I. The digestion-resistant bands correspond to the *rbbp4Δ4* allele. B Gels of rb1 exon 2 PCR amplicons. The 7bp deletion and wildtype alleles can be distinguished based on band size.

**S1 Table.** Related to Fig 3. Frequency of progeny genotypes and phenotypes from heterozygous *rbbpΔ4/+* crossed to heterozygous *rbbpΔ4/+;Tg*(*Tol2<ubi:rbbp4-2AGFP>*) zebrafish. Three independent crosses were set up between female *rbbpΔ4/+* heterozygotes and male *rbbpΔ4/+;Tg*(*Tol2<ubi:rbbp4-2AGFP>*) transgenic heterozygotes. Embryo clutches were sorted into positive and negative GFP groups at 2dpf. At 5dpf larvae were sorted by phenotype and then genotyped. “Total” column shows approximately 50% GFP- and GFP+ in each clutch, indicating the *Tg*(*Tol2<ubi:rbbp4-2AGFP>*) line contains a single transgene integration. Scoring of GFP negative embryos revealed close to Mendelian segregation of the *rbbpΔ4/Δ4* gross mutant phenotype in one quarter of the progeny (28.2%, 21.3%, 29%). A random sampling of embryos from clutches 2 and 3 were genotyped and confirmed Mendelian segregation of the phenotype with the *rbbpΔ4* allele. In each clutch, none of the GFP positive embryos showed the rbbp4 mutant phenotype. Random sampling and genotyping confirmed ~ one quarter of morphologically normal GFP+ larvae were homozygous mutant *rbbpΔ4/Δ4* (32.4%, 20.8%, 20.8%).

| Cross | Total # of progeny | GFP Negative |  |  |  |  |  | GFP Positive |  |  |  |  |  |
| --- | --- | --- | --- | --- | --- | --- | --- | --- | --- | --- | --- | --- | --- |
|  |  | Total GFP- | Normal Phenotype | Mutant Phenotype | *rbbp4* +/+ | *rbbp4* Δ4/+ | *rbbp4* Δ4/Δ4 | Total GFP+ | Normal Phenotype | Mutant Phenotype | *rbbp4* +/+ | *rbbp4* Δ4/+ | *rbbp4* Δ4/Δ4 |
| Clutch 1 | 76 | 39/76 (51.3%) | 28/39 (71.8%) | 11/39 (28.2%) | n/a | n/a | n/a | 37/76 (48.7%) | 37/37 (100%) | 0/37 (0%) | 8/37 (21.6%) | 17/37 (45.9%) | 12/37 (32.4%) |
| Clutch 2 | 108 | 61/108 (56.4%) | 48/61 (78.6%) | 13/61 (21.3%) | 5/24 (20.8%) | 11/24 (45.8%) | 8/24 (33.3%) | 47/108 (43.5%) | 47/47 (100%) | 0/47 (0%) | 8/24 (33.3%) | 11/24 (45.8%) | 5/24 (20.8%) |
| Clutch 3 | 307 | 155/307 (50.5%) | 110/155 (71%) | 45/155 (29%) | 18/48 (47.3%) | 17/48 (35.4%) | 13/48 (27%) | 152/307 (49.5%) | 152/152 (100%) | 0/152 (0%) | 16/48 (33.3%) | 22/48 (45.8%) | 10/48 (20.8%) |

**S2 Table.**
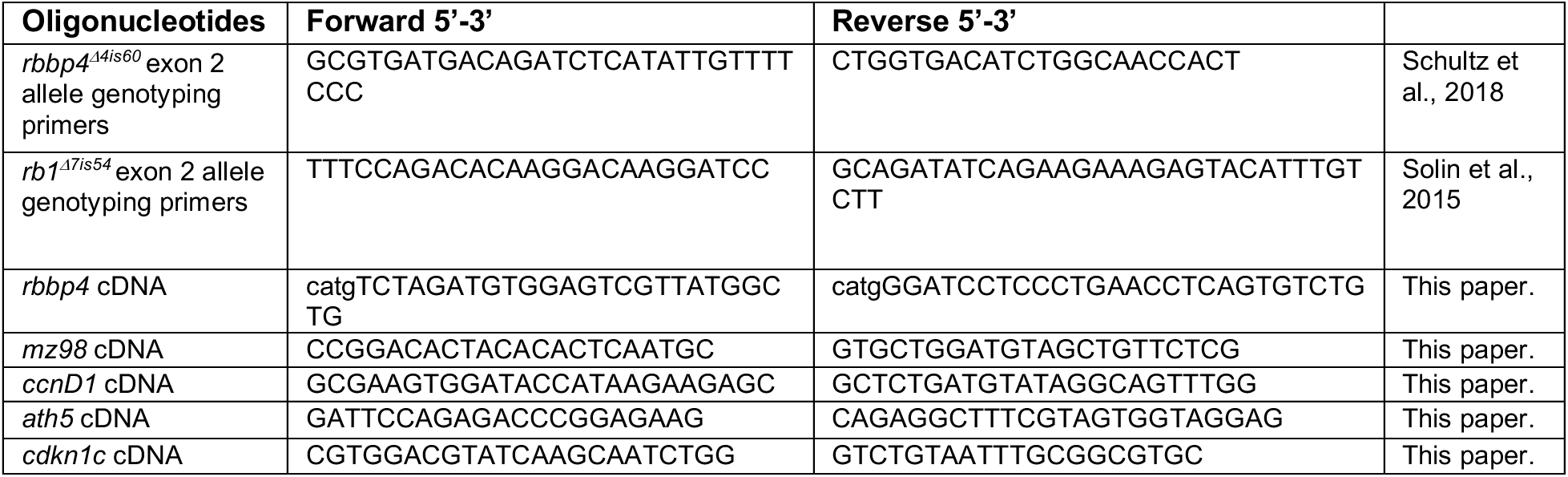
Sequences of primers used in this study.

